# Deletion of GPR39 Prevents Pulmonary Arterial Hypertension by Attenuating Hypoxia-Induced Aberrant Signaling

**DOI:** 10.64898/2026.06.27.735008

**Authors:** Carmen Methner, Lijuan Liu, Allura Thompson, Mary Plascencia, Priyanka Chakravarty, Sanjiv Kaul

## Abstract

Pulmonary arterial hypertension (PAH) is a devastating disease with poor outcome affecting relatively young subjects. The arachidonic acid (AA) metabolite, 15-hydroxyeicosatetraenoic acid (15-HETE), has been implicated in the pathogenesis of hypoxia-induced PAH. We tested the hypothesis that genetic deletion of GPR39, the target receptor for 15-HETE, will attenuate PAH. We subjected wild-type (WT) and GPR39 KO to 4 weeks of hypoxia versus normoxia, after which right ventricular and systemic hemodynamics were measured. Immunohistochemistry of lung was performed for pulmonary arteriolar thickness as well as capillary and pericyte density. Lung tissue was also analyzed for AA and 15-HETE levels as well as signaling events (mRNA and protein levels) downtream of GPR39 activation. Unlike WT mice, GPR39 KO mice did not develop PAH. They also exhibited markedly less pulmonary ateriolar remodeling and greater pulmonary capillary density. mRNA expression of genes in the Gα_q_, Gα_s_and Gα_12/13_ pathways were upregulated in the WT mice while GPR39 KO hypoxic showed no change in these genes. WT and not GPR39 KO hypoxic mice exhibited enhanced AKT phosphorylation. Downstream of the phosphatidylinositol 3-kinase-AKT pathway, endothelial nitric oxide synthetase was upregulated in both WT hypoxia and GPR39 KO hypoxia mice, while sonic hedgehog was upregulated only in WT hypoxia mice. We conclude that hypoxia-induced aberrant signaling is markedly attenuated with genetic deletion of GPR39, which is associated with less pulmonary arteriolar remodeling and greater capillary density, thus preventing PAH. These results suggest that pharmacological inhibition of GPR39 may offer a novel treatment for PAH.

## Introduction

Pulmonary arterial hypertension (PAH), albeit rare, is a devastating disease characterized by elevated pulmonary pressures resulting from pathologic pulmonary arteriole remodeling and vasoconstriction, ultimately resulting in right ventricular failure and death.^1^ Despite therapeutic advances including combination pulmonary vasodilator therapy and the more recent emergence of activin signaling inhibition^2,3^, the overall prognosis remains dire. Novel approaches that address mechanisms upstream of the known metabolic abnormalities in PAH have the potential for significantly altering management of these patients.

15-Hydroxyeicosatetraenoic acid (15-HETE), a metabolite of arachidonic acid (AA), has been implicated in the pathogenesis of hypoxia-induced PAH.^4^ 15-Lipoxygenase (15-LO) that converts AA to 15-HETE is upregulated in VSMCs and endothelium of patients with PAH.^5,6^ Plasma 15-HETE levels are elevated in animal models of hypoxia-induced PAH^4^ as well as in humans^7^ where they are associated with increased mortality.^8^ Mice fed 15-HETE develop PAH with its antecedent histopathological features.^9^

15-HETE contributes to PAH by altering several signaling pathways that are implicated in the pathobiology of PAH, including pulmonary vascular modeling and vasoconstriction. Specifically, 15-HETE inhibits apoptosis in pulmonary vascular smooth muscle cells (VSMCs)^10,11^, increases resting intracellular Ca^++^ ^12^, and inhibits K^+^ channels.^13–15^ It also stimulates adventitial fibrosis.^9,16^ The receptor through which 15-HETE acts was unknown until recently.

We have shown previously that 15-HETE serves as the endogenous agonist of the G-protein coupled receptor 39 (GPR39) in cardiac VSMCs and pericytes.^17,18^ GPR39 has significant constitutive activity via its Gα_q_ and Gα_12/13_ subunits.^19,20^ When stimulated by Zn^++^ or synthetic agonists, in addition to further activation of Gα_q_ and Gα_12/13,_ Gα_s_ is also activated.^19,20^ Gα_q_ activation by 15-HETE increases cytosolic Ca^++^ and causes extracellular signal related kinase (ERK) phosphorylation.^17^ Increased cytosolic Ca^++^ causes VSMC^17^ and pericyte contraction^18^, while phosphorylated ERK translocates to the nucleus and activates several transcription factors.^21,22^

Activated Gα_12/13_ stimulates the AKT-phosphatidylinositol 3-kinase (PI3K) pathway, which affects several nuclear transcriptional factors leading to cell growth and proliferation as well as reduced apoptosis.^23,24^ The AKT-PI3K pathway also acts as a regulator of hypoxia-induced factors 1 and 2 (HIF-1 and HIF-2)^25^, endothelial nitric oxide synthetase (eNOS)^26^, and cyclooxygenase 2 (COX2)^27^, and influences vascular endothelial growth factor A (VEGFA) regulation.^28^ HIF-1, in turn, binds to the endothelin-1 promotor causing formation of endothelin-1.^29^ The AKT-PI3K pathway also activates the sonic hedgehog (SHH) pathway.^30,31^

Activation of both Gα_q_ and Gα_12/13_stimulates the Rho pathway leading to cell proliferation and growth through, among other molecules, Rho-associated coiled-coil kinase (ROCK). Finally, Gα_s_ activation stimulates cyclic adenosine monophosphate response element binding protein (CREB), a kinase inducible transcription factor responsible for cell proliferation and growth. Since all these pathways are operative in PAH, we hypothesized that animals with genetic deletion of GPR39 will not develop 15-HETE-induced PAH compared to their wild-type (WT) littermate controls when subjected to hypoxia.

In this study we show, for the first time, that isolated primary lung VSMCs and pericytes express GPR39, which is also present in pulmonary arterioles and in whole lung tissue. Unlike their WT littermates, GPR39 knockout (KO) or heterozygous (HET) mice do not develop PAH or right ventricular (RV) dysfunction after 4 weeks of hypoxia, and pulmonary arterial remodeling is attenuated in these mice. Hypoxic GPR39 KO mice have significantly increased lung capillary density compared to WT hypoxia mice. Only WT hypoxia mice showed increased messenger ribonucleic acid (mRNA) expression of AKT1, CREB, ERK, and ROCK2, as well as phosphorylated AKT. Similarly, mRNA expression of signaling molecules associated with hypoxia further downstream was also higher in WT hypoxia mice and either unchanged or downregulated in GPR39 KO hypoxia mice. Apart from increased levels of eNOS and SHH, other protein levels were not increased in WT hypoxia mice. Taken together these results indicate that, in addition to less pulmonary arteriolar remodeling in hypoxic GPR39 KO mice, increased capillary density contributes to lower microvascular resistance, leading to the lower pulmonary artery pressure. These effects are associated with GPR39 deletion that attenuates mRNA expression of AKT1, CREB, ERK, and ROCK2 as well as common hypoxia-induced molecules in the Gα_q_, Gα_s_, and Gα_12/13_ pathways. Thus, pharmacological targeting of GPR39 may offer a novel therapeutic approach for treating PAH.

## Methods

All animal procedures performed in this study were approved by the Institutional Animal Care and Use Committee of Oregon Health & Science University and adhered to NIH Guide for the Care and Use of Laboratory Animals. All reporting is based on ARRIVE guidelines. One hundred and ten mice of both sexes were used. Our study design accounted for sex as a biological variable. Our study examined males and female and similar results were reported for both sexes, so they have been combined.

C57Bl6 mice of both sexes (12 to 24 weeks old) were used as WT. When GPR39 KO^17,18^ and HET mice (genotype confirmed by the polymerase chain reaction [PCR]) of both sexes were used, their WT littermates served as controls. Three groups of animals were used: Group 1 included WT and GPR39 KO mice for detection of GPR39 by immunocytochemistry (ICC) from isolated primary lung VSMCs and pericytes as well as immunohistochemistry (IHC) of pulmonary arterioles. They also underwent whole lung qPCR for GPR39 mRNA expression.

Groups 2 and 3 mice were randomized to either 4 weeks of hypoxia or normoxia. Group 2 mice underwent right heart hemodynamic measurements and lung immunohistochemistry (IHC) for vascular (arteriole, capillary, and capillary pericyte) changes. For Group 3 animals, lungs were sectioned into two. One section was used for mass spectroscopy, and the other section was used for western blotting and qPCR. Group 1 and 3 mice were anesthetized with 3% isoflurane prior to euthanasia with cervical dislocation prior to tissue removal. Tissue 15-HETE and AA levels were measured using mass spectroscopy. qPCR and western blotting was performed for AKT1, ERK 1/2, ROCK2, CREB, eNOS, iNOS, HIF-1_α_ a, HIF-2_α_, COX-2, VEGFA, prostaglandin-E (PGE), and endothelin-1, as well as the SHH, smoothened (SMO), suppressor of fused (SUFU) and glioma-associated Oncogene Homolog 1 (GLI1). qPCR was also performed for prostaglandin-I (PGI).

### Cell Isolation and culture

VSMCs and pericytes were isolated from 6 WT and 6 GPR39 KO mice (Table 1A) using modifications of our previously published approaches.^32^ For VSMC isolation, lung slices were placed into a collagen (#C5533, Sigma-Aldrich, St. Louis MO)-coated tissue culture plate, 1 slice per well of a 24-well plate, containing 100 µL fetal buffered saline (FBS). The plate was placed in a cell culture incubator (37 °C; 95% air and 5% CO2) for 4 h after which 500 µL of VSMC culture medium was added to each well. The medium was changed 5 days later and incubated for a further 7 days, allowing for migration of VSMCs from the lung slices onto the tissue culture plastic. The lung slices were removed from the wells and discarded. Cells were enzymatically detached from wells using trypsin: ethylenediaminetetraacetic acid (EDTA, 0.05%: 0.5 M). Once cells had detached, 0.2 mL Dulbecco’s modified Eagle’s medium (DMEM)+10% FBS per well was used to inactivate the enzyme. The cell suspension was transferred to a 15 mL conical tube, pooled, and centrifuged at 1000 rpm for 8 min. The supernatant was removed, and cells were re-suspended in 10 mL of smooth muscle cell culture medium.

**Table 1A.** Group 1 WT Animals Used for GPR39 Identification by ICC, IHC and qPCR (n=16)

| Genotype | Number | Age (weeks mean ± SEM) | Sex (M, F) |
| --- | --- | --- | --- |
| WT | 8 | 13.4 ± 0.6 | (4, 4) |
| KO | 8 | 14.0 ± 0.4 | (4, 4) |

**Table 1B.** Group 2 Animals Undergoing Hypoxia or Normoxia Used for Hemodynamics and IHC for Pulmonary Artery Thickness and Capillary Density Measurements(n=74)

| Genotype | Number | Age (weeks mean ± SEM) | Sex (M, F) | Treatment |
| --- | --- | --- | --- | --- |
| WT | 7 | 13.4 ± 0.6 | (5, 2) | Normoxia |
| WT | 13 | 14.0 ± 0.4 | (5, 8) | Hypoxia |
| GPR39-HET | 14 | 12.0 ± 0.5 | (7, 7) | Normoxia |
| GPR39-HET | 8 | 14.6 ± 0.7 | (5, 3) | Hypoxia |
| GPR39-KO | 15 | 11.9 ± 0.6 | (12, 3) | Normoxia |
| GPR39-KO | 17 | 15.1 ± 0.5 | (8, 9) | Hypoxia |

Primary lung pericytes were isolated by dicing lung tissue and digesting with collagenase (#LS004176, CLS-2, Worthington, Lakewood, NJ,) in an agitated water bath (37°, 100 RPM) for 70 min. This mixture was triturated every 20 min using a 6″ long 14-gauge metal cannula attached to a 35 mL disposable syringe. After the final trituration, the suspension was passed through a 70 µm disposable cell strainer and collected in a 50 mL conical tube. Additionally, 10 mL DMEM was passed through the cell strainer to collect any remaining cells. The cell suspension was centrifuged at 1000 rpm for 10 min at room temperature. The cells were resuspended in 6 mL DMEM + 60 µL CD31 (#553369, BD Pharmingen, Franklin Lakes, NJ) conjugated Dynabeads Sheep anti-Rat IgG (#11035, Invitrogen, Carlsbad, CA) in a 15 mL conical tube and placed on a rotator (medium setting) for 40 min at room temperature. The tube was mounted into a magnetic separator for 1 min allowing the beads (with cells attached) to adhere to the sides. The supernatant containing unbound cells without platelet endothelial adhesion molecule 1 (CD31-) was collected and further isolated with platelet derived growth factor β (PDGFRβ, CD140b. #14-1402-82, Invitrogen) using the same method saving the bound PDGFRβ cells. The resulting pericytes (CD31- & PDGFRβ+) were placed in a T75 collagen coated flask and grown until confluence.

### Immunocytochemistry

ICC for primary lung cells was performed using our recently published methods.^32^ VSMCs and pericytes placed on glass coverslips, were fixed in fresh 4% paraformaldehyde in phosphate buffered saline (PBS: 0.1 M sodium phosphate buffer, 0.9% NaCl, pH 7.4) and subsequently blocked with 5% goat serum in PBS containing 0.3% Triton X-100 and 1% Bovine Serum Albumin for 90 min at room temperature, then incubated overnight at 4 °C with primary antibodies diluted in blocking buffer. The following primary antibodies and dilutions were used: rabbit anti-GPR39, 1:100, (#ABIN1048812, Antibody online, Limerick, PA) rabbit anti-NG2, 1:100 (#AB5320, Millipore Sigma< Burlington, MA) and mouse anti-α-SMA (#A2547, Millipore Sigma,). Cells were washed with PBS + 0.1% Tween 20 and secondary antibody (1:200, Alexa 488-conjugated goat anti-rabbit or Alexa 594-conjugated donkey anti-rabbit, and Alexa 647-conjugated donkey anti-mouse (#A11008, #A21207 and # A31571, respectively. They were placed in blocking buffer for 2 h at room temperature. Cell nuclei were labeled with Hoechst 33342 (#62249, Thermo Fisher). The coverslips were washed and mounted using Fluoromount-G (#0100-01, Southern Biotech, Birmingham, AL). Images were acquired with a confocal microscope (Nikon Eclipse Tie-A1RSi, Tokyo, Japan).

### Immunohistochemistry

For IHC of pulmonary arterioles and capillaries tissue was fixed in fresh 4% paraformaldehyde in PBS. Hydrated and deparaffinized sections were placed in 10 mM sodium citrate buffer (pH 6.0) at a sub-boiling temperature for 30 min. After cooling the slides, a 30 min antigen retrieval was initiated. The slides were washed in PBS and then immersed in PBS containing 4% normal goat serum, 1% bovine serum albumin, and 0.3% Triton X-100 for 90 min at room temperature.

The sections were then incubated overnight at 4 °C with primary antibodies: 1:300x, Lectin I (GSL I, Isolectin B4, Bionylated, # B-1205, Vector, Newark, CA; 1:100 α-smooth muscle (α-SMA, #NB300978, Novus, Littleton, CO). Secondary antibodies for Lectin and α-SMA were cyanine 3 (Cy3) streptavidin (1:800x,#434315 Invitrogen) and 6474-Donkey anti Goat (H+L 1:400x #A21147). 4’-6-diamidino 2 phenylimadole (DAPI. 1:6000x, # H3570, Invitrogen) was then used to stain the nuclei. Pericytes were labeled with neuroglial antigen 2 (NG2). Sections were incubated with rabbit anti-NG2 antibody (1:100, #AB5320, Millipore Sigma), and donkey-anti-rabbit IgG Alexa Fluor 594 (1:400, #A21207, Invitrogen). After coverslips were placed, images were acquired with a confocal microscope (Nikon Eclipse Tie-A1RSi). For measuring PA wall thickness, images were acquired at 60x magnification and for measuring capillary and pericyte density, 40x magnification was used. Three images acquired from each animal were analyzed in a blinded manner.

For GPR39 labeling, using the same protocol as above, sections were incubated with the following antibodies: rabbit anti-GPR39 antibody (1:200, #ABIN1048812, antibodies-online), Biotinylated Griffonia Simplicifolia Lectin 1 (1:300, #B-1205, Vector labs), Cy3-streptavidin (1:800, #434315, Invitrogen) and anti-rabbit IgG Alexa Fluor 488 (1:400, #A21206, Invitrogen).

### qPCR

qPCR was performed to determine the relative gene expression within whole lung tissue, which was extracted after euthanasia and lungs immediately flash-frozen in liquid nitrogen. RNA quality was inspected and verified on a NanoDrop prior to further use.

The relative amount of mRNA for each gene was calculated by utilizing the Comparative Ct method (ΔΔCt) normalized to β-actin as an endogenous control. ΔΔCt values were then used to find the relative fold change in expression (2^(-(ΔΔCt))) of each group in relation to the expression in the whole wildtype normoxia lung tissue.

Approximately 500 ng of RNA was reverse-transcribed to generate first-strand complementary DNA (cDNA) with a High-Capacity cDNA Reverse Transcription Kit with RNAse inhibitor (# 4374966, Thermo Fisher). cDNA was subjected to PCR amplification using gene-specific Taqman primers (Thermo Fisher Scientific). The 10 μL reaction containing Taqman Fast Advanced Master Mix (# 4444557, Thermo Fisher), Taqman primers (GPR39: Mm07298841_m1; β Actin: Mm05794665_g1, iNOS: Mm00440502_m1; eNOS: Mm00435217_m1; COX2: Mm00478374_m1; Endothelin-1: Mm00438659_m1; HIF1α: Mm00468869_m1; HIF2α: Mm01236112_m1; Prostaglandin I_2_: Mm00801939_m1; SHH: Mm00436528_m1; SUFU: Mm00489385_m1; SMO: Mm01162710_m1; GLI1: Mm00494654_m1; VEGFA: Mm00437306_m1; CREB: Mm00501607_m1, ROCK2: Mm01270843_m1; ERK: Mm00442479_m1; AKT1: Mm01331626_m1; Prostaglandin E: Mm00460181_m1, Thermo Fisher); RNA, and nuclease free H2O. The experiments were performed by following Taqman’s instruction with Quantstudio 7 Flex Real-Time PCR System (RRID:SCR_022651, Applied Biosystems, Waltham, MA).

### Western Blot

Lung tissue was homogenized in liquid nitrogen and lysed in Ripa lysis buffer (#PI89900, Thermo Fisher) with Pierce protease and phosphatase inhibitor (#A32959, Thermo Fisher). The homogenates were centrifuged at 14000 rpm for 20 min at 4°C. The Pierce BCA Protein assay kit (#23225, Thermo Fisher) was used to measure the concentration of protein lysate in samples. Protein (20-30 µg) per lane were separated on 4%–12% SDS-polyacrylamide gels and transferred to polyvinylidene difluoride membranes. Total protein staining was performed using Licor Bio Revert® 700 total Protein Stain (#926-11016, Licor, Lincoln, NE) following the manufacturer’s protocol. The membranes were blocked with Licor Intercept® TBS blocking buffer (#927-60003, Licor) for 60 min followed by primary antibodies blocking buffer with 0.02% Tween-20: 1:1,000 rabbit anti-SHH antibody (Cell Signaling, Cat. #2207), 1:2,000 rabbit anti-SMO antibody (Thermo Fisher Scientific, Cat. #PA5-113312), 1:1,000 rabbit anti-SUFU antibody (Cell Signaling, Cat. #2522), 1:1,000 mouse anti-GLI1 antibody (Proteintech, Cat. #66905-1), 1:1,000 rabbit anti-VEGFA antibody (abcam, Cat. #AB46154), 1:3,000 rabbit anti-Endothelin 1 antibody (abcam, Cat. #AB117757), 1:1,000 rabbit anti-Hif-1 alpha antibody (Novus biologicals, Cat. #NB100-479), 1:1,000 rabbit anti-Hif-2 alpha antibody (Novus biologicals, Cat. #NB100-122),1:1,000 mouse anti-eNOS antibody (Cell Signaling, Cat. #5880), 1:1,000 mouse anti-iNOS antibody (Novus biologicals, Cat. #NB222119), 1:500 mouse anti-COX2 antibody (Thermo Fisher Scientific, Cat. #35-8200), 1:200 rabbit anti-Prostaglandin E Synthase antibody (Cayman, Cat. #160140), 1:120 mouse anti-Prostaglandin I Synthase antibody (Cayman, Cat. #100023), 1:1,000 rabbit anti-CREB antibody (Cell Signaling, Cat. #4820), 1:1,000 rabbit anti-Phospho-CREB antibody (Cell Signaling, Cat. #9198), 1:2,000 mouse anti-ERK1/2 antibody (Cell Signaling, Cat. #4696), 1:2,000 rabbit anti-Phospho-ERK1/2 antibody (Cell Signaling, Cat. #4370), 1:1,000 rabbit anti-AKT antibody (Cell Signaling, Cat. #9272), 1:1,000 rabbit anti-Phospho-AKT antibody (Cell Signaling, Cat. #9271), 1:1,000 rabbit anti-ROCK2 antibody (Cell Signaling, Cat. #8236). After 1h incubation with secondary antibody (IRDye® 800CW Goat anti-Rabbit IgG Secondary Antibody, 1:5000, Licor Bio, Cat. #926-32211, IRDye® 800CW Goat anti-Mouse IgG Secondary Antibody, 1:5000, Licor Bio, Cat. #926-68070) at room temperature, the membranes were imaged using Li-Cor Odyssey Imager with Image Studio software. The intensity of each protein band was normalized to total protein stain in each sample using Empiria Studio 2.2 software (Li-Cor Inc, Lincoln, NE).

### Mass Spectroscopy

Lung tissue from Group 2 mice (Table 1C) were used for mass spectroscopic analysis for 15-HETE and AA. Lungs were recovered after euthanasia. Working solutions and internal standards were prepared in 2:1 chloroform:methanol. Lung tissue was homogenized with commercial bead beater tubes with 1.4 mm ceramic beads in methanol:water (66:34 v/v) at 100 mg/mL for 30 s at 5m/s with antioxidant mix consisting of 0.2 mg/mL each of butylated hydroxytoluene, and ethylenediaminetetraacetic acid, and 2 mg/mL each of triphenyl phosphine and indomethacin at 10 µL/mL. Lung tissue homogenate was then placed at -80°C for 1 hour, spun to pellet insoluble debris for 15 s at 13,000 RCF.

**Table 1C.**
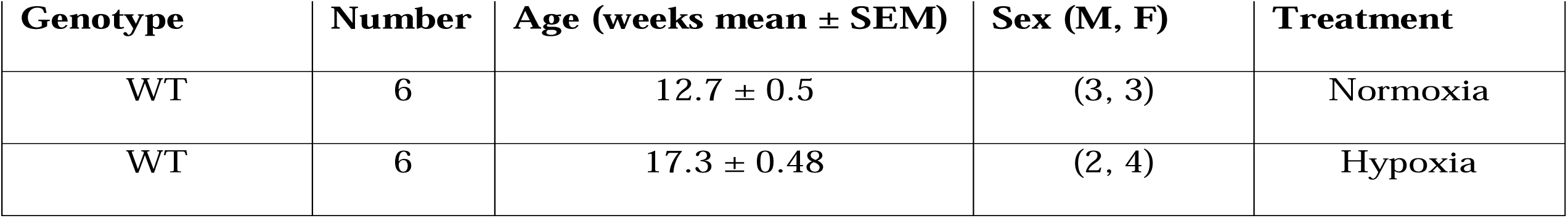
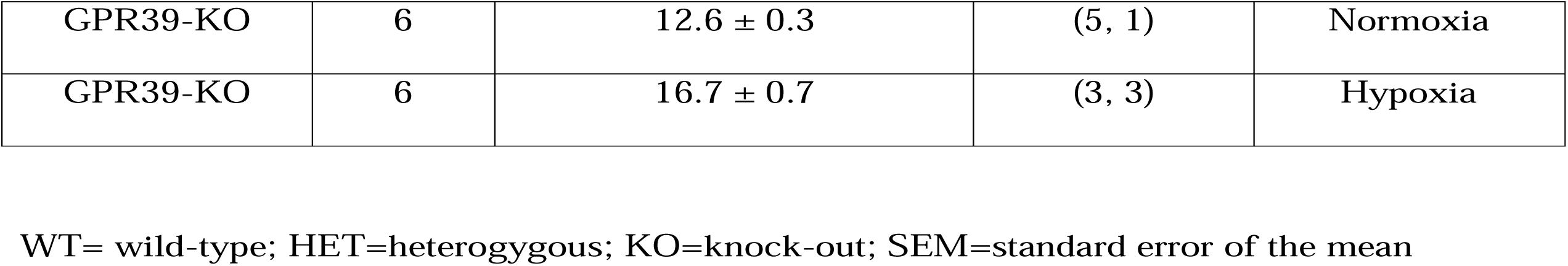
Group 3 Animals Undergoing Hypoxia or Normoxia Used for Lung Tissue Metabolomics, qPCR, and Western Blot (n=24)

| Genotype | Number | Age (weeks mean ± SEM) | Sex (M, F) | Treatment |
| --- | --- | --- | --- | --- |
| WT | 6 | 12.7 ± 0.5 | (3, 3) | Normoxia |
| WT | 6 | 17.3 ± 0.48 | (2, 4) | Hypoxia |
| GPR39-KO | 6 | $12.6 \pm 0.3$ | (5, 1) | Normoxia |
| GPR39-KO | 6 | $16.7 \pm 0.7$ | (3, 3) | Hypoxia |

WT= wild-type; HET=heterozygous; KO=knock-out; SEM=standard error of the mean

For 15-HETE measurement samples were loaded by gravity onto Oasis HLB 60 mg pre-conditioned with 1 mL each of methanol and 0.1% formic acid in water as previously described.^33^ After initial slow gravity load, 1-2 psi positive pressure nitrogen was used to finish load. Samples were washed with 2×1 mL 10% methanol 0.1% formic acid at 1-2 psi. Following wash, cartridges were dried for 5 minutes at high positive pressure to remove all water. Eluent was collected from cartridges in 13×100 collection tubes using elution with 1×1 mL 0.1% formic acid in methanol, followed by 1×1 mL of ethyl acetate. A trap solution, 20% glycerol in methanol (20 mL) was added to the top of the combined eluates and samples were subsequently dried at 40°C 1 L/minute for 45 min using a Biotage Turbovap unit (Uppsala, Sweden). Samples were resuspended in 20 mL of methanol, vortexed, followed by addition of 80 mL of 0.1% formic acid in 80:20 water: acetonitrile. Samples were filtered through 0.2 µ bioinert Eppendorf filter, transferred to vials and analyzed. An unextracted curve was used across the range 10-1000 pg/sample.

15-HETE was measured using MS/MS with a 5500 Q-TRAP hybrid/triple quadrupole linear ion trap mass spectrometer (Applied Biosystems) with electrospray ionization (ESI) in negative mode. The mass spectrometer (MS/MS) was interfaced to a Shimadzu (Columbia, MD) SIL-20AC XR auto-sampler followed by 2 LC-20AD XR LC pumps and analysis on an Applied Biosystems/SCIEX Q5500 instrument. The instrument was operated with the following settings: source voltage -4000 kV, GS1 70, GS2 70, CUR 45, TEM 750 and CAD gas MED. The gradient mobile phase was delivered at a flow rate of 0.5 mL/min, and consisted of two solvents, A: 0.05% acetic acid in water and B: acetonitrile. The LC column used was a Kinetex Phenyl-Hexyl 50×2.1 mm 2.6 µM. The initial concentration of solvent B used was 20%, this was held for 0.5 minutes, increased to 60% B over 10.5 minutes, increased to 95% B over 1 min, held at 95% B for 1 min, and then returned to initial conditions over 0.1 min, followed by re-equilibration for 1.9 min for a total run time of 15 min. The column oven was set to 50°C, autosampler to 15°C with needle rinse before and after each injection. Data was acquired using Analyst 1.7.1, and analyzed with Multiquant 3.03.

For AA, we modified a previously described method.^34^ Tissue slurry sample (50 μL), reference standard and internal standard were mixed in screw-capped glass tubes and 2 mL of methylation mixture was added (methanol/acetyl chloride, 20:1 v/v). The mixture was incubated at 25 °C with shaking at 100 rpm for 12 hours. After incubation, methyl esters were isolated using liquid-liquid extraction performed by adding 0.5 mL of water and 1.0 mL of hexane to each sample tube, except for heart effluent which was extracted with 0.25 mL of hexane to concentrate the sample. The mixture was vortexed for 20 s and centrifuged for 5 min at 2000 rpm. The upper (organic) layer was removed and placed into auto-sampler vial for analysis using gas chromatography-mass spectrometry (GC-MS).

Analysis was performed using a 7890B GC with a 5977A MS detector with an autosampler and injector (Agilent, Santa Clara, CA) operated in split mode. The column was an Agilent DB-FastFAME column (30 m, 0.25 mm id, 0.25 μm film thickness). Helium was the carrier gas at a flow rate of 1 mL/min. The injection port and auxiliary heater were maintained at 245°C. A 1 μL sample was injected in split mode (1:10) at an initial oven temperature of 150°C, held for 0.5 min and then increased at 15°C/min to 200°C, followed by 25°C/min to 240°C, held for 2 min and then returned to 50°C. The MS was operated at a source temperature of 230°C and a MS quad temperature of 150°C in positive electron impact mode. AA was detected at *m/*z 43,55, 67, 74, 79, and 91, each with a 25 msec dwell time after a solvent delay of 4 min. The retention time for AA was 7.7 min. The GC-MS was controlled and data acquired using enhanced MassHunter software version B.07.04.2260 and results analyzed using Agilent MassHunter quantitative analysis software version B.07.0. Calibration curves for quantification were generated from peak area ratios for authentic standard:internal standard for calibrator samples.

### Hypoxia Exposure

Animals were placed in a hypoxia chamber with a ProOx 110 O_2_ controller (Biospherix, Parish, NY, USA). Chamber O_2_ was maintained at 10%. The chamber was opened once a week to refresh cages and water bottles and to top up food. The animals experienced the same light cycle as their normoxia littermate controls that were held in animal housing (room air, 21% O_2_).

### Measurement of Right Heart Hemodynamics

Mice were anesthetized (3% induction and 1.5% maintenance), intubated, and connected to a volume-control ventilator (Harvard apparatus; Holliston, MA, USA). Mice were placed in supine position on a warming pad to maintain a body temperature at 37°C.

A PE20 catheter was placed in the right femoral vein for administration of fluids as needed. A PE10 catheter was inserted in the left femoral artery and connected to AxonCNS Digidata 1440A interfaced with AxoScope 10.7 (Molecular Devices, San Jose, CA, USA) for continuous monitoring and recording of mean aortic pressure. The right internal jugular vein was exposed and a SPR-1000 Mikro-Tip mouse pressure catheter (Millar Instruments, Houston, TX, USA) interfaced to the AxonCNS Digidata 1440A system was advanced through a small incision into the right ventricle (RV) with continuous pressure monitoring to confirm the correct location. This catheter has a 1F catheter tip size, a 20 cm usable catheter length, and a 0.8F catheter body.

Hemodynamic measurements were performed at 1, 3, and 5 min from continuous cardiac cycles after placement of the Millar catheter. These included heart rate, mean aortic pressure, peak RV systolic pressure (RVSP), and RV end-diastolic pressure. Heart rate was measured from the mean interval between the peak RV systolic pressures from 10 consecutive beats. Maximum RV diameter and RV wall thickness were measured from postmortem sections of the heart at the mid-papillary muscle level and RV wall stress (dynes/cm^2^) was calculated using the formula: (P·r)/2h, where P = RVSP, r = RV radius, and h = RV wall thickness.

At the end of the experiment, while maintaining an airway pressure of 15 mm Hg, the animal was perfused at systemic pressures with 4% formaldehyde in PBS. Mice were euthanized by cervical dislocation, the lungs removed for IHC and the heart removed for measuring RV dimensions.

### Statistical Methods

Power calculation was performed for sample size estimation where hypoxia was estimated to result in a 30% increase in RVSP compared to normoxia in WT mice. Unless otherwise noted, all data are expressed as mean ± 1SEM. Comparisons between groups were evaluated by ANOVA followed by Sidak post hoc test. Differences were considered significant at *p* < 0.05.

All data presented in this paper will be available from the senior author upon request.

## Results

### Group 1 animals

The demographics of animals used for detection of GPR39 by ICC, IHC, and qPCR are shown in Table 1A. Figure 1A illustrates ICC results from isolated primary lung VSMCs for GPR39. DAPI, α-SMA, and GPR39 staining are shown along with merged images. Figure 1B shows ICC of a primary lung pericyte showing DAPI, GPR39, α-SMA and NG2 staining. Since the antibodies for GPR39 and NG2 are of rabbit origin, these represent images from different animals. Both VSMCs and pericytes clearly express GPR39.

**Figure 1:**
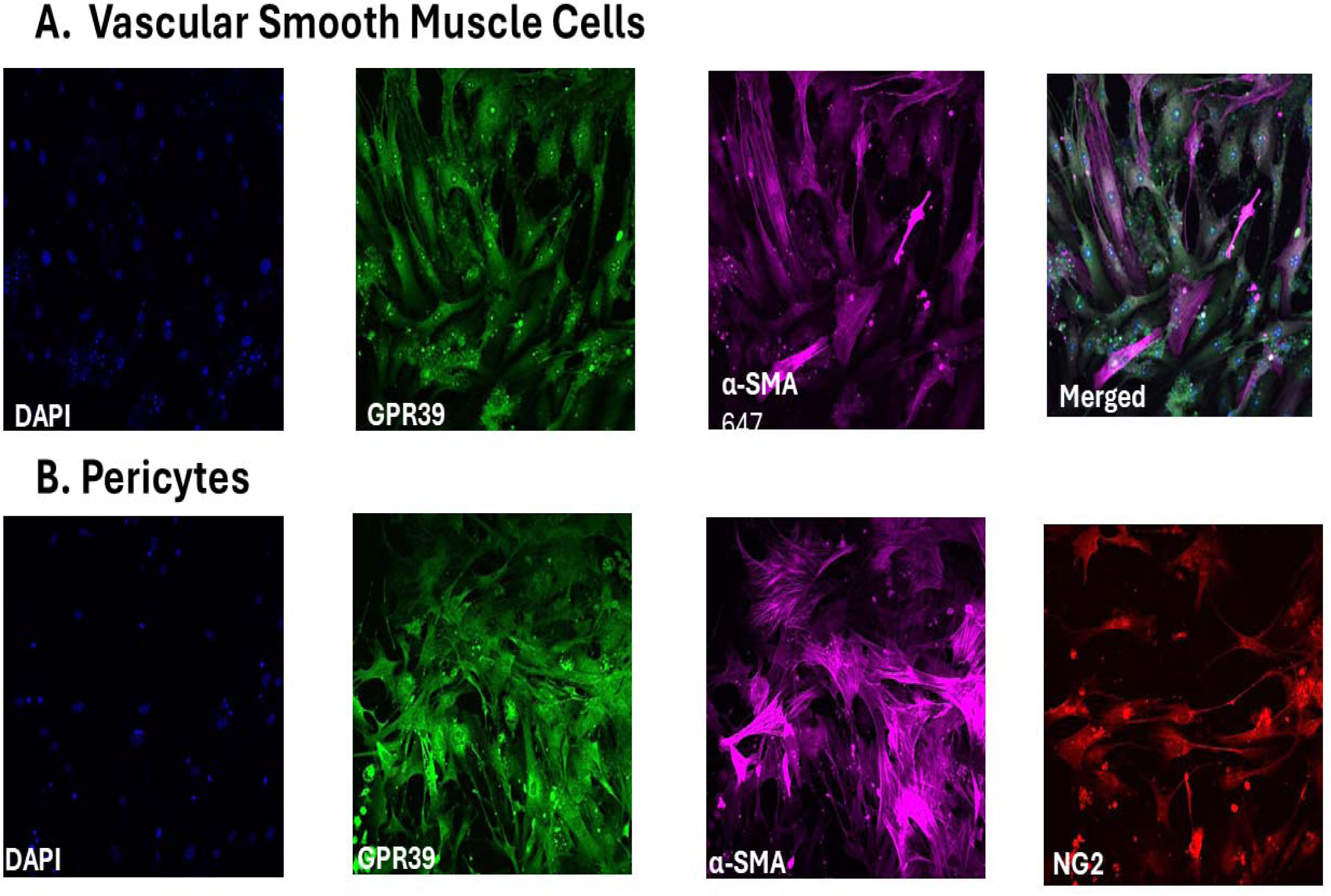
ICC results from Group 1 animals. (A) primary lung VSMCs staining for DAPI, GPR39, and α-SMA staining along with merged images are shown. (B) primary lung pericyte staining for DAPI, GPR39, α-SMA and NG2. Since the antibodies for GPR39 and NG2 are of rabbit origin, these represent images from different animals. Both VSMCs and pericytes clearly express GPR39.

Figure 2A illustrates IHC of lung tissue for presence of GPR39 in pulmonary arterioles from a WT mouse and Figure 2B shows the same from A GPR39 KO mouse. Antibodies for DAPI, α-SMA, and GPR39 were used, and a merged image is also shown. There is abundant presence of GPR39 in the pulmonary arterioles of WT but not GPR39 KO mouse. Figure 2C depicts results of qPCR for GPR39 gene expression in whole lung tissue from WT and GPR39 KO mice. As expected, lungs from WT mice express GPR39 while lungs from KO mice do not.

**Figure 2:**
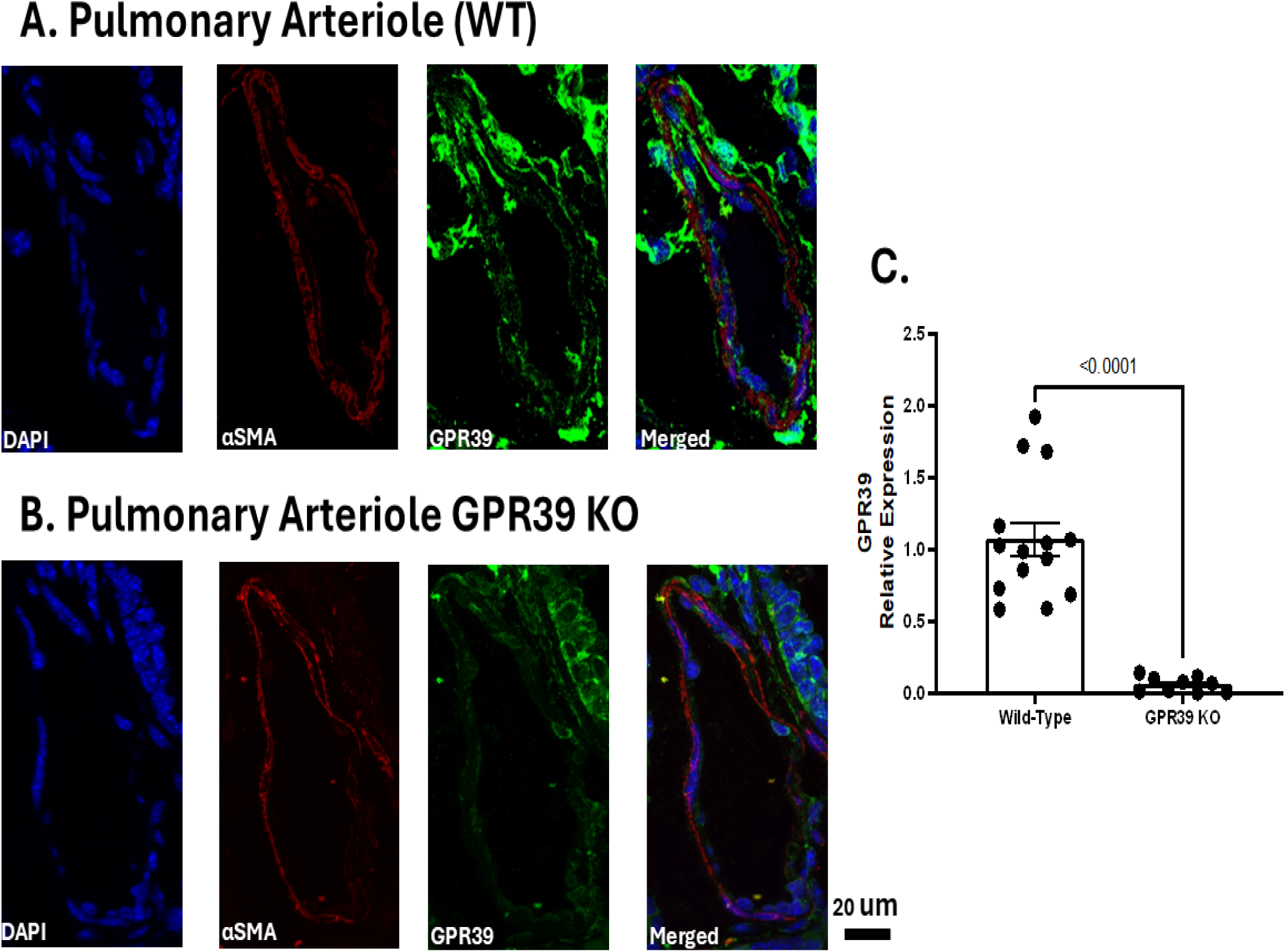
IHC and qPCR results from Groups 1 animals. IHC of lung tissue for presence of GPR39 in VSMCs from a WT (A) and GPR39 KO (B) mouse. Antibodies for DAPI, α-SMA, and GPR39 were used, and a merged images are also shown. There is abundant presence of GPR39 in the pulmonary arterioles of WT but not GPR39 KO mouse. Panel C shows results of qPCR for GPR39 expression in lung tissue from WT and GPR39 KO mice. As expected, lungs from WT mice express GPR39 while lungs from KO mice do not.

### Group 2 animals

These 74 animals underwent hemodynamic measurements followed by IHC of the pulmonary vasculature. Table 1B lists the number, genotypes, sex, and treatment (hypoxia versus normoxia) of these animals. In 4 animals RV pressures could not be attained because of aberrant venous anatomy.

#### Hemodynamic results

Figure 3 illustrates the RV hemodynamic results from the Group 2 animals. Panels A and B are examples of pressure tracings from WT normoxia and WT hypoxia mouse, respectively. The normoxia mouse exhibits normal RVSP and RVEDP, while in the hypoxia mouse both are elevated, with characteristic systolic notching associated with poor pulmonary circulation compliance.^35^ Panels C and D are examples of pressure tracings from a GPR39 KO normoxia and a GPR39 KO hypoxia mouse, respectively. RV pressures in the GPR39 KO normoxia mouse are similar to that of the WT normoxia mouse, while the hypoxia GPR39 KO mouse shows slightly elevated right ventricular systolic (RVSP) but not right ventricular end-diastolic pressure (RVEDP) compared to the normoxia mouse. Both RVSP and RVEDP in the GPR39 KO mouse are markedly less than in the WT hypoxia mouse. There was no difference between GPR39 heterozygous and GPR39 KO mice exposed to normoxia or hypoxia (data not shown). Panels E shows aggregate results from the 71 Group 2 animals for RVSP, RVEDP, and RV wall stress, respectively. In all instances these variables are significantly elevated in the WT hypoxia mice compared to both WT normoxia as well as GPR39 KO hypoxia mice. Table 2 depicts average heart rate and mean aortic pressure derived from 3 measurements taken at 1,3, and 5 min after insertion of right heart catheter from all animals. There is no difference between heart rate and mean aortic pressure between the 6 groups of animals.

**Figure 3:**
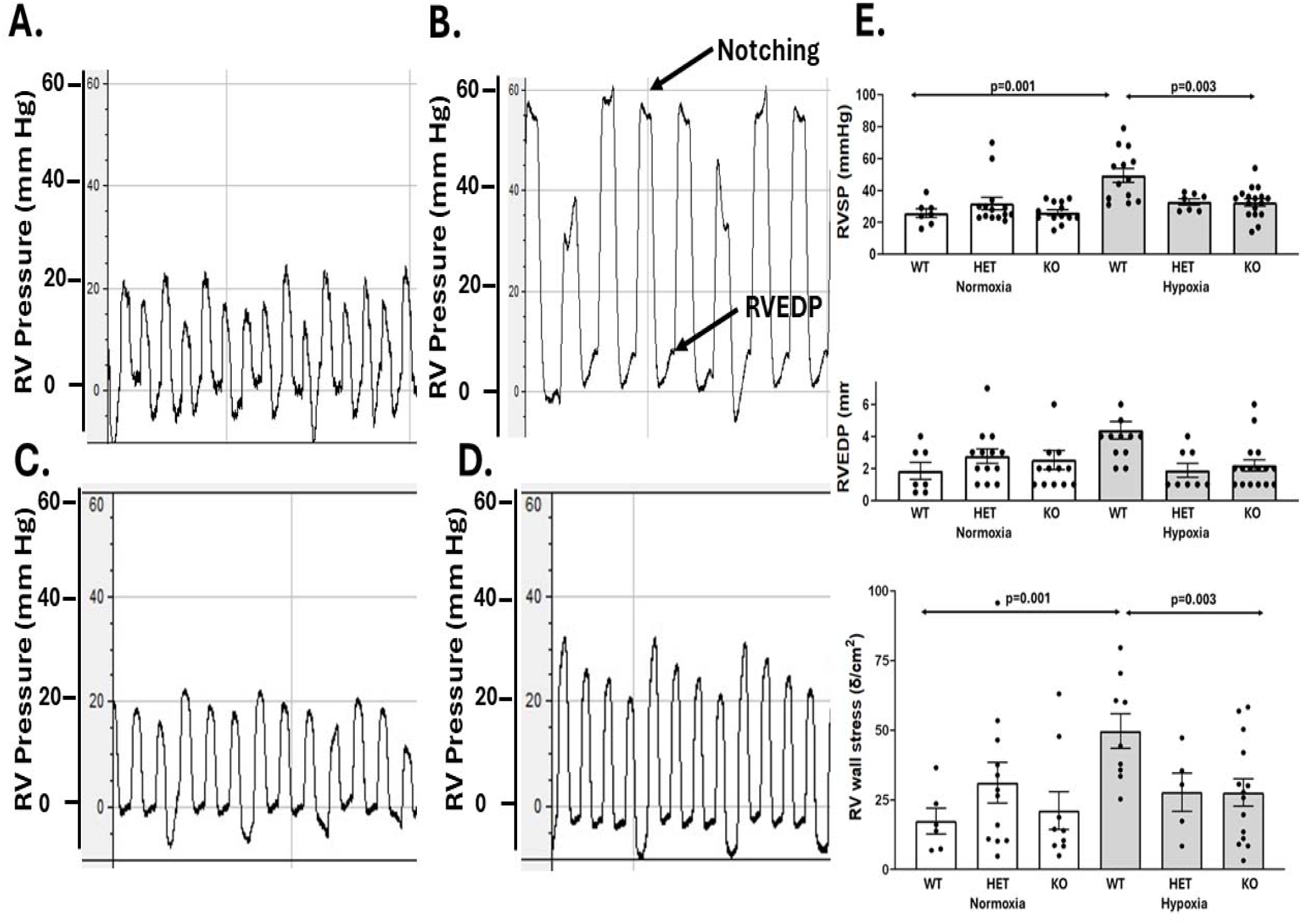
Right ventricular hemodynamics from Group 2 animals. Panels A and B are examples of pressure tracings from a WT normoxia and a hypoxia mouse, respectively. The normoxia mouse exhibits normal RVSP and RVEDP, while in the hypoxia mouse both are elevated, with characteristic systolic notching associated with poor pulmonary circulation compliance.^35^ Panels C and D are examples of pressure tracings from a GPR39 KO normoxia and a GPR39 KO hypoxia mouse, respectively. RV pressures in the GPR39 KO normoxia mouse are similar to that of the WT normoxia mouse, whereas the hypoxia GPR39 KO mouse shows slightly elevated RVSP but not RVEDP. There was no difference between GPR39 heterozygous and GPR39 KO mice exposed to normoxia or hypoxia (data not shown). Panels E shows the aggregate results from 71 Group 2 animals for RVSP, RV-end-diastolic pressure, and RV wall stress, respectively. In all instances these variables are significantly elevated in the WT hypoxia mice compared to both WT normoxia as well as GPR39 KO hypoxia mice.

**Table 2.** Heart Rate and Mean Aortic Pressure in Group 2 animals (mean±1SEM)

| **Genotype** | **Number** | **Treatment** | **Heart Rate**<br>**(beats/min)** | **Mean Aortic**<br>**Pressure (mm Hg)** |
| --- | --- | --- | --- | --- |
| WT | 7 | Normoxia | 445 ± 61 | 55 ± 6 |
| WT | 13 | Hypoxia | 473 ± 44 | 69 ± 2 |
| **P-value** |  |  | >0.999 | 0.238 |
| HET | 14 | Normoxia | 502 ± 13 | 61 ± 2 |
| HET | 7 | Hypoxia | 438±67 | 68 ± 3 |
| **P-value** |  |  | 0.908 | 0.877 |
| KO | 13 | Normoxia | 498 ± 24 | 52 ± 4 |
| KO | 17 | Hypoxia | 508 ± 32 | 65 ± 4 |
| **P-value** |  |  | 0.998 | 0.172 |

p-values two way ANOVA

#### Immunohistochemistry for Pulmonary Arteriolar Wall Thickness

Figure 4 illustrates IHC of lung pulmonary arterioles stained for DAPI, α-SMA, and lectin at 60x from a WT normoxia (upper panel A) and a WT hypoxia (upper panel B) Group 2 animal. PA wall is significantly thicker in the hypoxia animal. Lower Panels A and B show examples form normoxia and hypoxia GPR39 KO animals, respectively. While the PA wall is thicker in the hypoxia GPR39 KO compared to the normoxia GPR39 KO animal, it is not as thick as in the WT hypoxia animal. The aggregate results from the Group 2 animals are shown in panel C. PA wall thickness is greater in both groups of hypoxia animals compared to littermate normoxia animals, but WT hypoxia animals have significantly thicker PA walls than hypoxia GPR39 KO animals.

**Figure 4:**
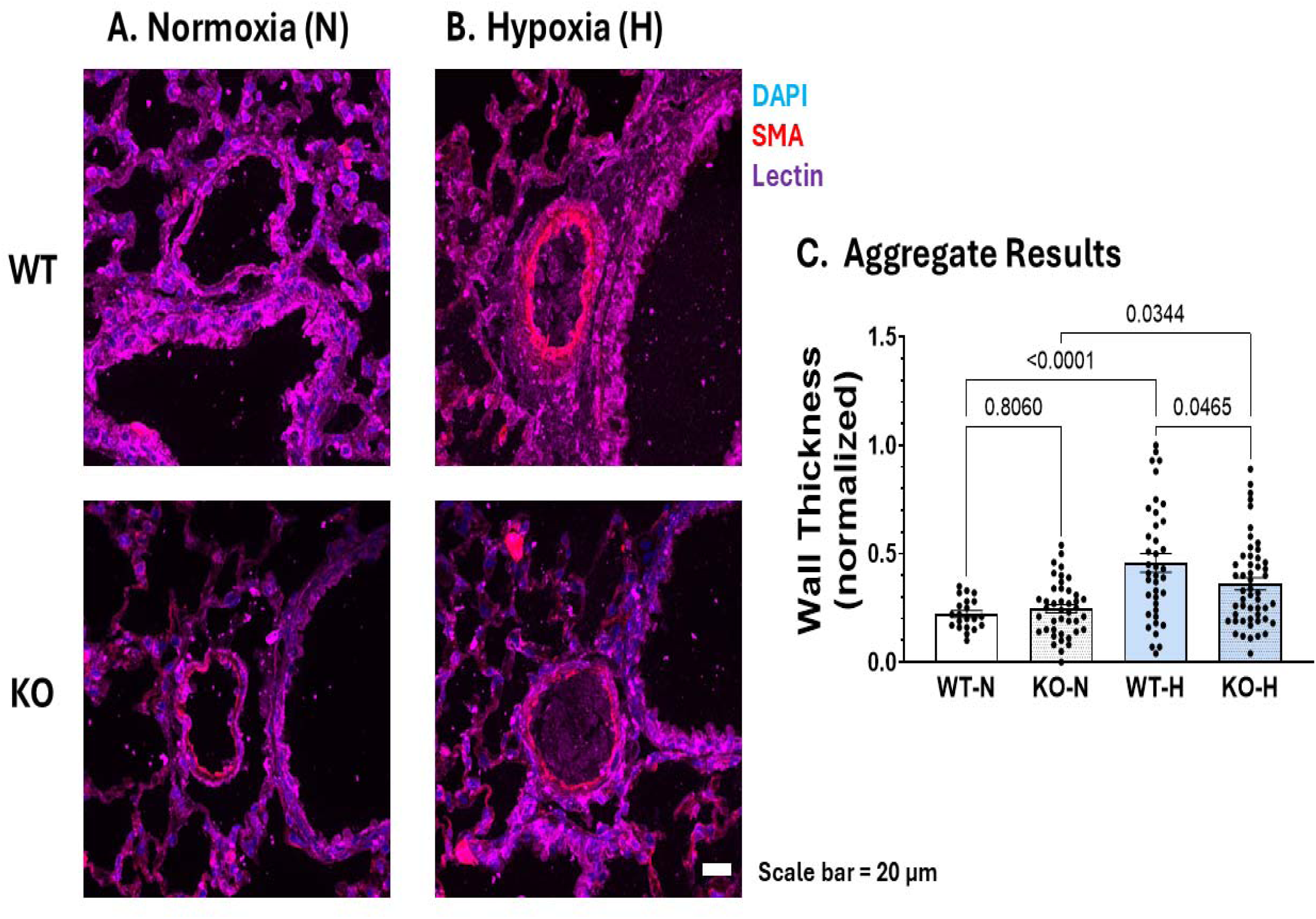
IHC results of pulmonary arterioles. Examples of lung tissue for DAPI, α-SMA, and lectin at 60x from Group 2 WT normoxia (upper panel A) and hypoxia (upper panel B) animals imaged at 60x. PA wall is significantly thicker in the hypoxia animal. Lower Panels A and B show examples form normoxia and hypoxia GPR39 KO animals. While the PA wall is thicker in the hypoxia GPR39 KO compared to the normoxia GPR39 KO animal, it is not as thick as in the WT hypoxia animal. Three different parts of the lungs were imaged in each animal, so each data point represents one image. The aggregate results are shown in panel C. PA wall is thicker in both groups of hypoxia animals compared to littermate normoxia animals, but WT hypoxia animals have significantly thicker PA walls than GPR39 KO animals.

#### Immunohistochemistry for Capillary Density

Figure 5 depicts examples of capillary density from the Group 2 WT and GPR39 KO mice during normoxia (top 2 panels) and hypoxia (bottom 2 panels), respectively. Images were acquired using channels specific for DAPI (panel A), lung tissue showing autofluorescence (panel B), and lectin for capillaries (panel C). Merged images are shown in panel D. Capillary density was expressed as a percent of total lung tissue. Staining for lectin is greatest in the GPR39 KO hypoxia mouse (panels C and D) compared to other groups. The aggregate results (panel E) show that capillary density in GPR39 KO hypoxia mice is increased compared to WT hypoxia and normoxia mice. However, there is no difference in the capillary density between hypoxia and normoxia GPR39 KO mice.

**Figure 5:**
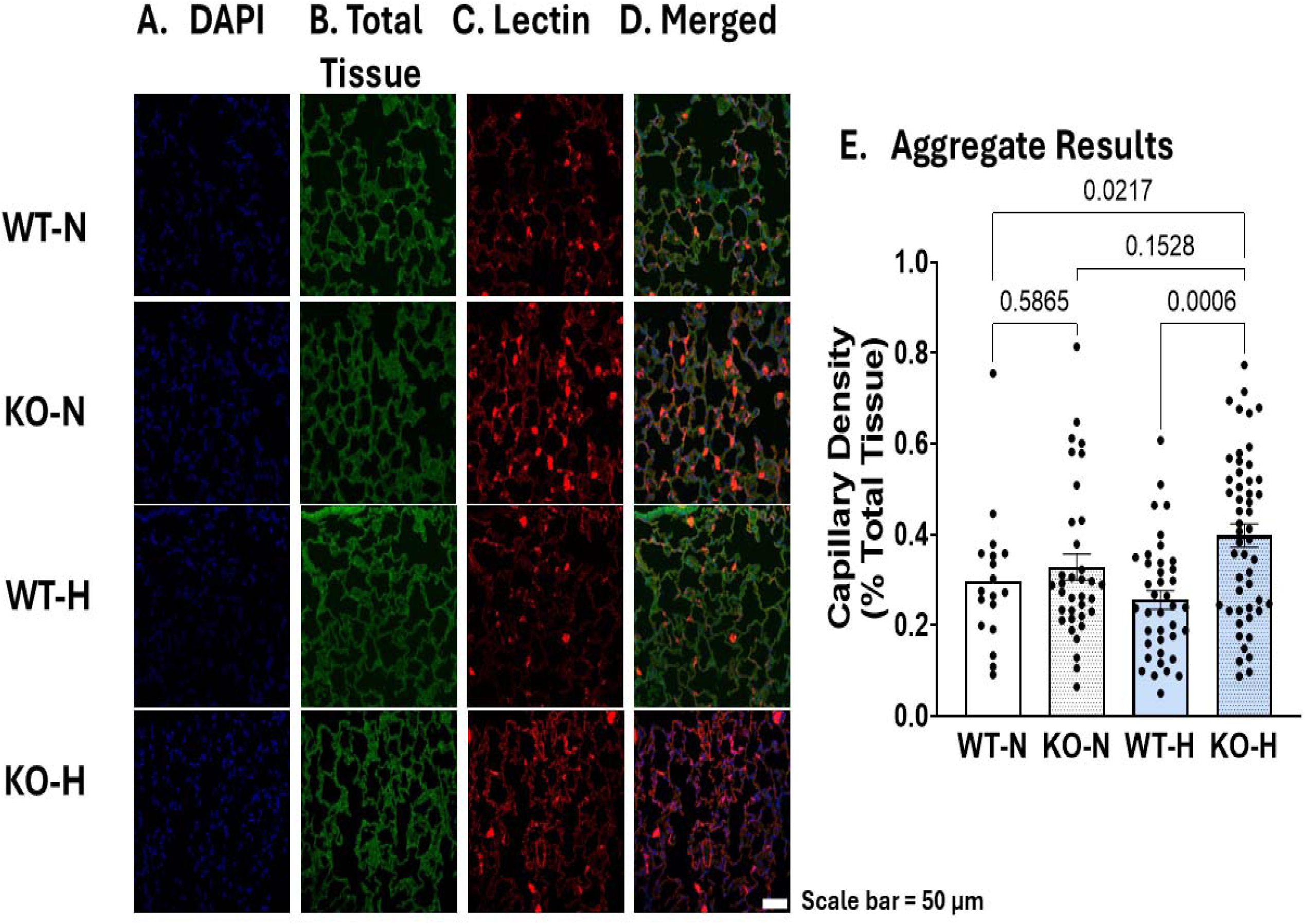
IHC Capillary density results. Examples from Group 2 WT and GPR39 KO during normoxia (top 2 panels) and hypoxia (bottom 2 panels) imaged at 40x. Images were acquired using channels specific for DAPI (panel A), background tissue showing autofluorescence (panel B), and lectin (panel C). Merged images are shown in panel D. Capillary density was expressed as a percent of lung tissue. From panels C and D it is evident that staining for lectin greatest in the GPR39 KO hypoxia tissue compared to other groups. Three different parts of the lungs were imaged in each animal, so each data point represents one image. The aggregate results (top panel E) show that capillary density in GPR39 KO hypoxia mice is increased compared to WT hypoxia and normoxia mice.

#### Immunohistochemistry for Capillary Pericyte Density

Figure 6 shows examples of pericyte distribution at the capillary level from Group 2 WT and GPR39 KO animals during normoxia (top 2 panels) and hypoxia (bottom 2 panels), respectively, imaged at 40x. Images were acquired using channels specific for DAPI (panel A), lung tissue exhibiting autofluorescence (panel B), and NG2 for pericytes (panel C). Merged images are shown in panel D. Pericyte density was expressed as percent of lung tissue. From panels C and D it is evident that there are more pericytes in both the normoxia and hypoxia GPR39 KO mice compared with normoxia or hypoxia WT mice. The aggregate results (panel E, top) show that capillary density in both normoxia and hypoxia GPR39 KO mice is increased compared to normoxia or hypoxia WT mice. Moreover, normoxia GPR39 KO mice have more pericytes than hypoxia GPR39 KO mice. The ratio of the pericyte covered area versus capillary covered area was similar between WT and GPR39 KO hypoxia animals (panel E, bottom panel) and this ratio in hypoxia GPR39 KO mice was less than in normoxia GPR39 KO mice, akin to that noted for pericyte density.

**Figure 6:**
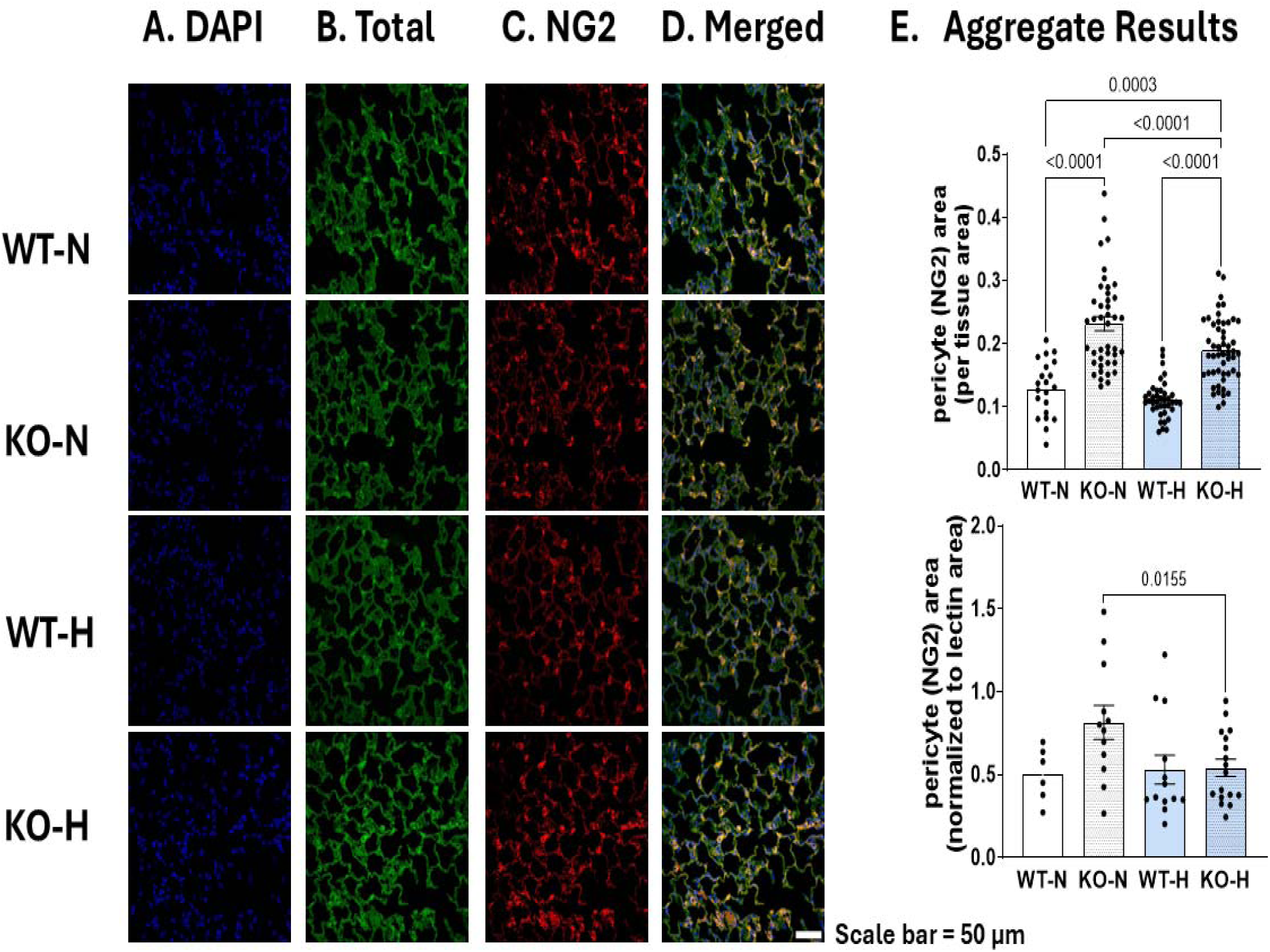
IHC Pericyte distribution at capillary level. Examples of pericyte distribution at the capillary level from Group 2 WT and GPR39 KO during normoxia (top 2 panels) and hypoxia (bottom 2 panels) imaged at 40x. Images were acquired using channels specific for DAPI (panel A), background tissue showing autofluorescence (panel B), and NG2 (panel C). Merged images are shown in panel D. Pericyte density was expressed as a percent of lung tissue. From panels C and D it is evident that NG2 shows more pericytes in the GPR39 KO tissue compared to WT mice and that GPR39 KO hypoxia mice had lower number of pericytes than WT hypoxia mice. Three different parts of the lungs were imaged in each animal, so each data point represents one image. The aggregate results (top panel E) show that capillary density in GPR39 KO mice is increased compared to WT mice. The pericyte/capillary ratio shows is normalized in hypoxia GPR39 KO mice and is lower than the normoxia GPR39 KO mice (bottom panel E).

### Group 3 animals

These 24 animals underwent either 4 weeks of hypoxia (n=12) or 4 weeks of normoxia (n=12), after which their lungs were harvested for mass spectroscopic measurements of 15-HETE and AA as well as IHC and qPCR. Table 1C lists the genotyping and sexes of these animals.

#### Mass Spectroscopy Results

Figure 7 shows mass spectroscopy results from lung tissue of normoxia (n=12) and hypoxia (n=12) animals. The WT and GPR39 KO animals are shown in different colors. Lung tissue AA levels are increased in hypoxia animals (panel A) as is the 15-HETE level (panel B). However, the ratio of 15-HETE/AA is similar in both groups (panel C) indicating that the elevated 15-HETE could result from increased substrate availability in addition to 15-LO upregulation, as previously reported.^4,5^

**Figure 7:**
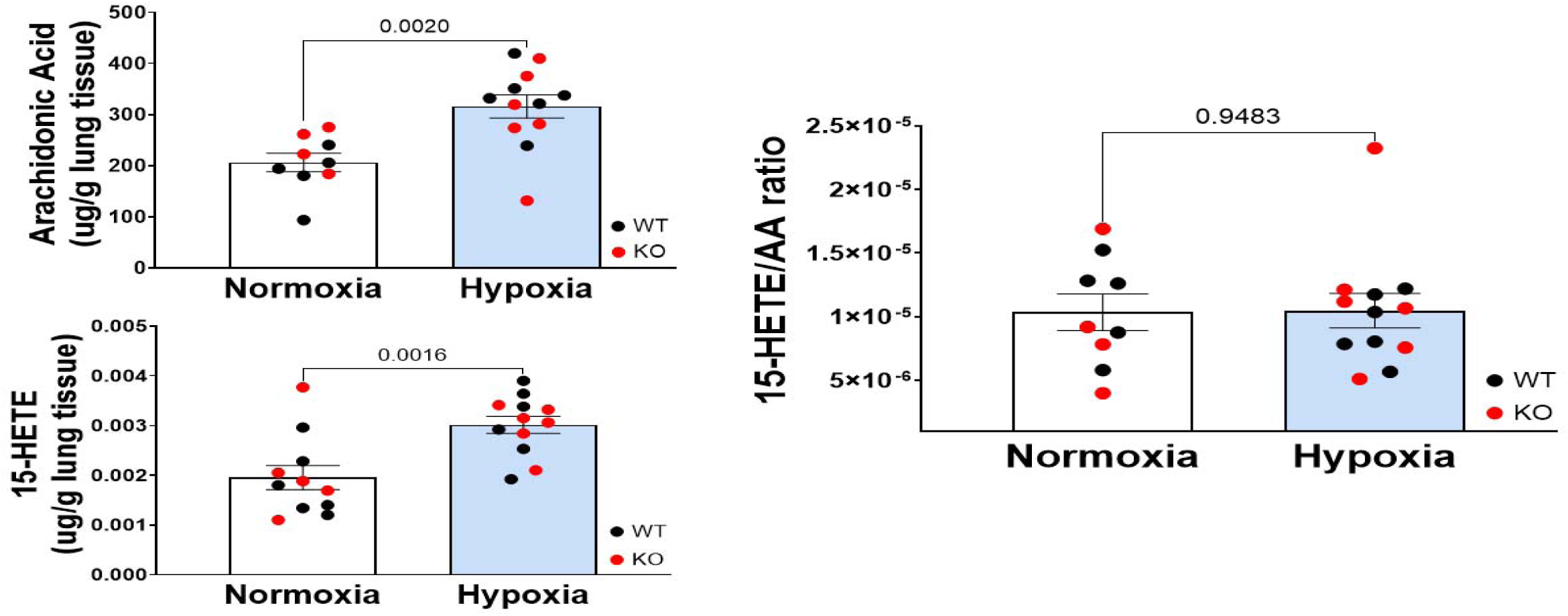
Mass spectroscopy results. AA (panel A) and 15-HETE (panel B) levels from normoxia (n=12) and hypoxia (n=12) animals, along with the 15-HETE/AA ratio (panel C). These animals were exposed to 4 weeks of either hypoxia or normoxia and lungs were harvested for measurement of AA and 15-HETE levels by mass spectroscopy. Each data point represents one animal. Lung tissue AA levels are increased in hypoxia animals as is the 15-HETE level. However, the ratio of 15-HETE/AA is similar in both groups. There were no differences between wild-type (black) and GPR39 knockout (red) mice and so their results are presented together.

#### mRNA results

Figure 8 depicts lung mRNA levels from the Group 3 animals. Each panel depicts aggregate results from the 4 groups of animals. All mRNA levels are significantly higher in the hypoxia WT mice compared to hypoxia GPR39 KO mice except for endothelin-1, where the difference did not reach statistical significance. The first column in Table 3 summarizes these results, which are also depicted in the central illustration.

**Figure 8:**
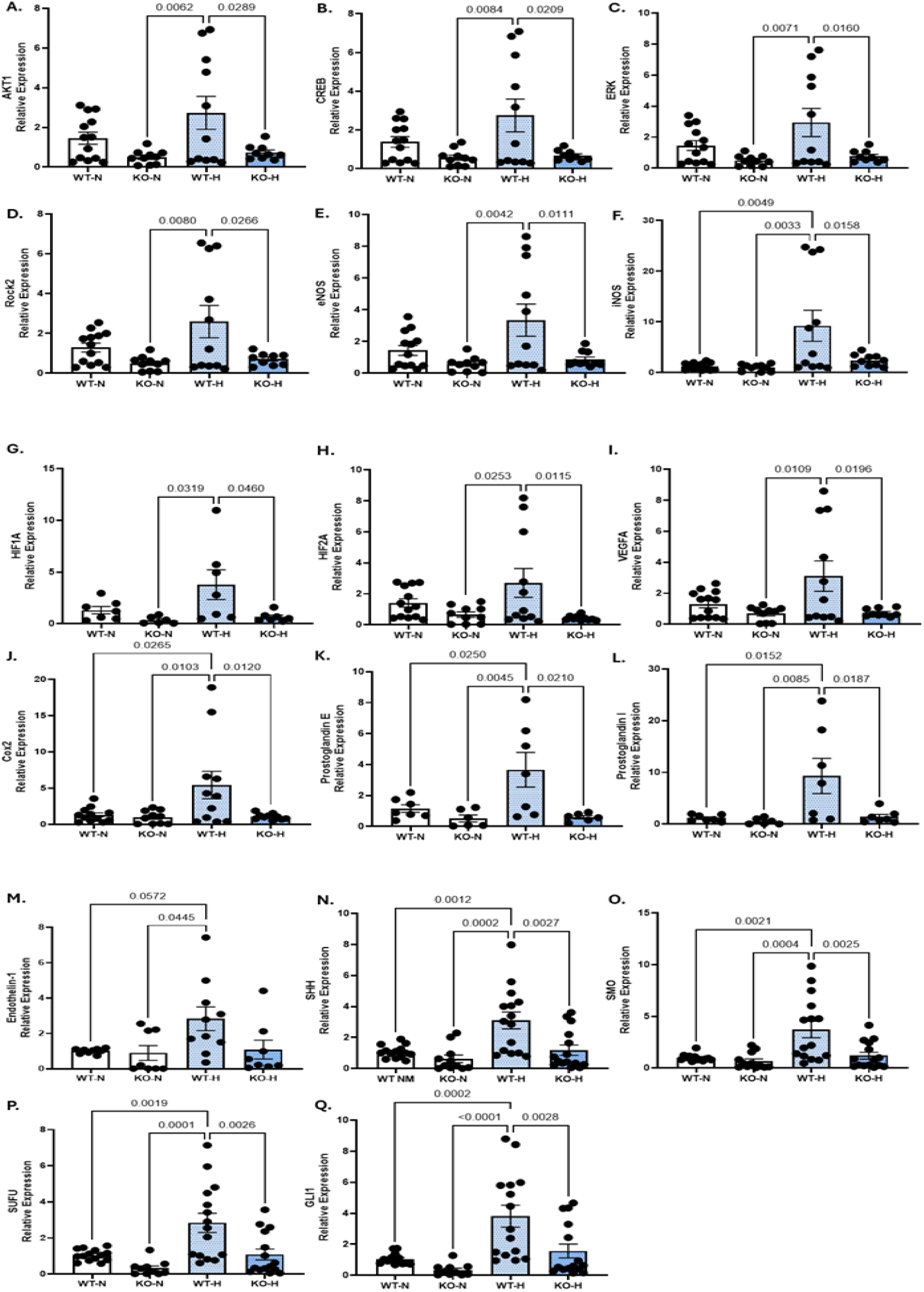
mRNA levels from the Group 3 animals for: (A) AKT, (B) CREB, (C) ERK, and (D) ROCK2, (E) eNOS, (F) iNOS, (G), HIF-1_α_, (H) HIF-2_α_, (I) VEGF, (J) COX-2, (K) prostaglandin E, (L) Prostaglandin I, (M) endothelin 1, (N) SHH, (O), SMO, (P) SUFU, and (Q) GLI1. Except for mRNA levels of endothelin-1, mRNA levels were increased in hypoxia WT compared to hypoxia GPR39 KO mice.

**Table 3.** Changes in mRNA Levels in Hypoxia WT Compared to GPR39 KO Mice And Changes in Total Protein Levels in Hypoxia Compared to Normoxia Mice (n=24)

|  | mRNA | Total Protein |  |
| --- | --- | --- | --- |
|  | WT vs. KO | WT | GPR39 KO |
| **AKT1** | Increased | Decreased | Decreased |
| **p-AKT1** |  | Increased | No change |
| **CREB** | Increased | Decreased | Decreased |
| **p-CREB** |  | Decreased | Decreased |
| **ERK 1 /2** | Increased | No change | No change |
| **p ERK 1 /2** |  | No change | No change |
| **ROCK2** | Increased | No change | No change |
| **eNOS** | Increased | Increased | Increased |
| **iNOS** | Increased | No change | No change |
| **HIF-1α** | Increased | Decreased | Decreased |
| **HIF-2α** | Increased | Decreased | Decreased |
| **VEGFA** | Increased | No change | No change |
| **COX2** | Increased | No change | No change |
| **Prostaglandin E** | Increased | No change | No change |
| **Prostaglandin I** | Increased |  |  |
| **Endothelin-1** | No change | Decreased | Decreased |
| **SHH** | Increased | Increased | No change |
| **SMO** | Increased | Decreased | Decreased |
| **SUFU** | Increased | Decreased | Decreased |
| **GL1** | Increased | Decreased | Decreased |

p=phosphorylated

#### Western Blot results

Figure 9 illustrates total and phosphorylated levels of AKT, CREB, ERK 1/2 and total level of ROCK2 from Group 3 animals. In each panel, under aggregate results for each of the 4 groups is an example of a western blot. AKT levels (panel A) were reduced in both groups of hypoxia mice compared to the normoxia mice. Phosphorylated AKT, normalized to total protein (panel B) as well total AKT (panel C) was, however elevated in the WT hypoxia and not the GPR39 KO mice compared to their normoxia controls. CREB levels normalized to total protein levels were markedly reduced in WT and GPR39 KO hypoxia mice compared to their normoxia controls (panel D). Phosphorylated CREB levels normalized to total protein (panel D) and total CREB (panel F) were almost undetectable in hypoxia mice compared to their normoxia controls. Total and phosphorylated ERK1/2 levels did not differ between the animal groups (panels G, H, I), neither did ROCK2 levels (panel J).

**Figure 9:**
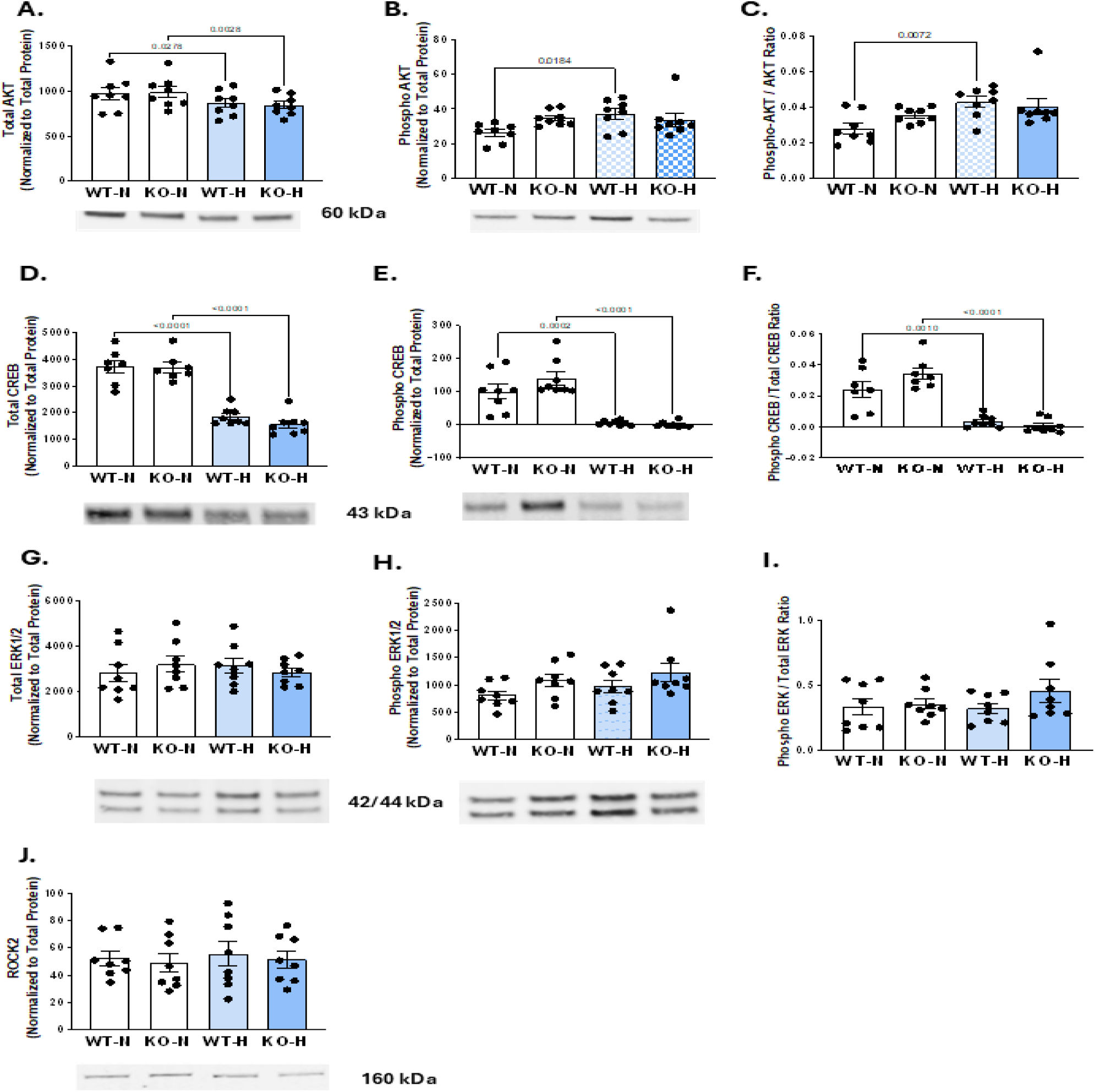
Total and phosphorylated protein levels from Group 3 animals. In each panel, under aggregate results for each of the 4 groups is an example of a western blot. Total AKT levels normalized to total protein levels are decreased with hypoxia in both WT and GPR39 KO mice (panel A). However, phosphorylated AKT levels normalized to total protein (panel B) or to total AKT (panel C) are elevated only in WT hypoxia mice compared to normoxia WT mice. Total CREB levels normalized to total protein levels are decreased in both hypoxia WT and hypoxia GPR39 KO mice compared to their normoxia controls (panel D). Similar results are seen for phosphorylated CREB whether normalized to total protein (panel E) or to total CREB (panel F). Total ERK 1/2 levels normalized to total protein levels are unaffected by hypoxia in both WT and GPR39 KO animals (panel G). Similar results are seen for phosphorylated ERK 1/2 whether normalized to total protein (panel H) or to total ERK 1/2 (panel I). ROCK2 levels normalized to total protein levels were also unaffected by hypoxia in both WT and GPR39 KO mice (panel J).

Figure 10 illustrates additional lung protein levels from the Group 3 animals. In each panel, under aggregate results for each of the 4 groups is an example of a western blot. eNOS expression is increased in both hypoxia groups compared to their normoxia controls (panel A), while there is no difference in iNOS expression between any of the groups (panel B). In contrast, HIF-1α and HIF-2_α_ were significantly downregulated in both WT and GPR39 KO groups with no difference between them (panel C). VEGFA and COX2 expressions were similar between all groups of anils (panels E and F). Endothelin-1 levels were decreased in hypoxic compared to normoxic mice, while PG2 levels were unchanged by hypoxia (data not shown). SHH levels were increased in WT and not GPR39KO hypoxia mice (panel G). In contrast SMO, SUFU, and GLI1) levels were significantly decreased in both WT and GPR39 KO hypoxic mice (panels H to J) with no differences noted within the hypoxia or normoxia groups. Columns 2 and 3 in Table 3 summarize the results of changes in proteins expression in both hypoxia groups compared to their normoxia controls, which is also depicted in the central illustration.

**Figure 10:**
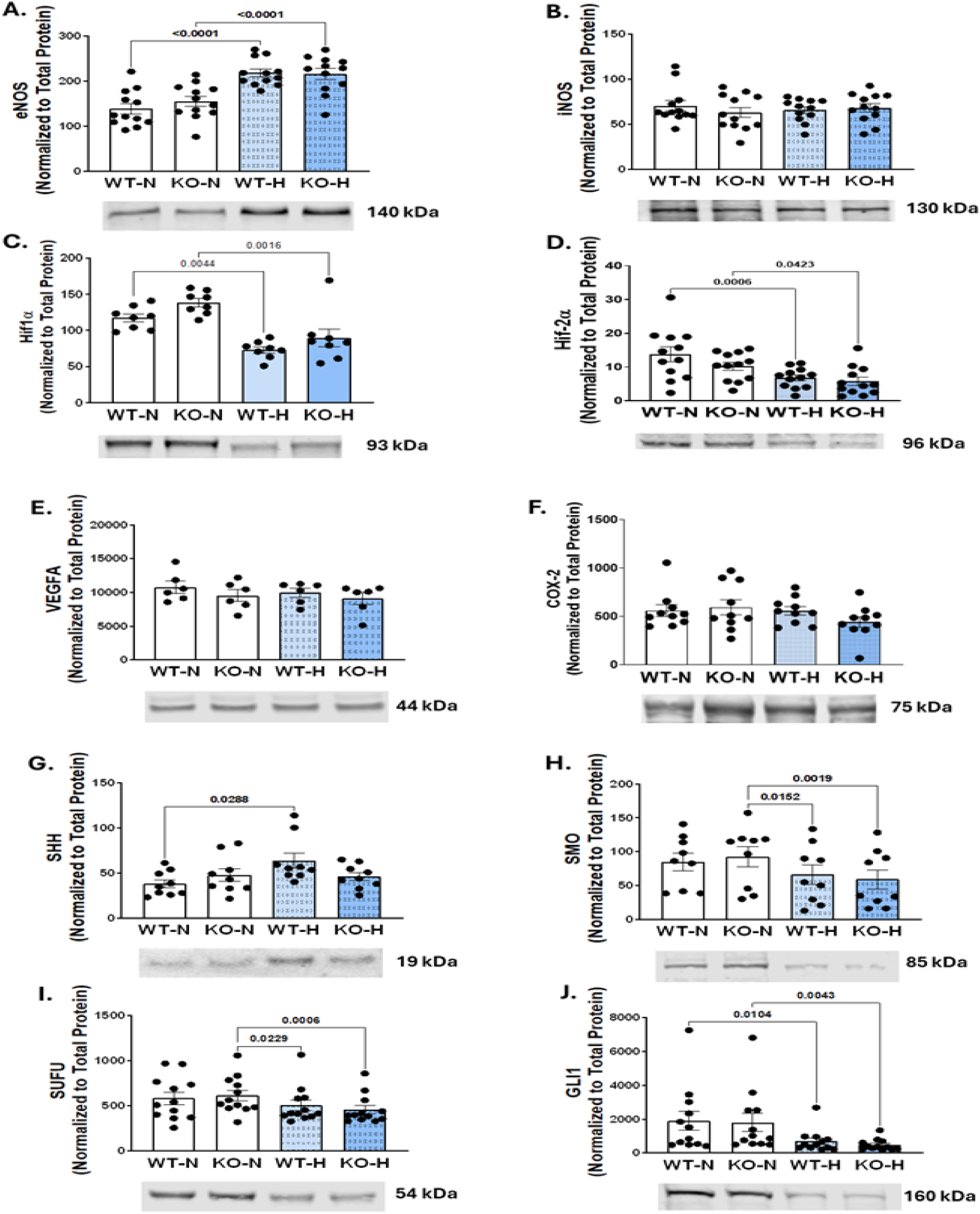
Protein levels from Group 3 animals. In each panel, under aggregate results for each of the 4 groups is an example of a western blot. eNOS expression was upregulated in both WT hypoxia and GPR39 hypoxia mice (panel A), while iNOS levels were similar between all four animal groups (panel B). HIF-1_α_ and HIF-2α were significantly downregulated in both WT and GPR39 KO hypoxia mice (panels C and D). In comparison, neither VEGFA nor COX2 expression was affected by hypoxia in WT or GPR39 KO mice (panels E and F). SHH levels were increased in WT and not GPR39KO hypoxia mice (panel G). In contrast SMO, SUFU, and GLI1 levels were significantly decreased in both WT and GPR39 KO mice (panels H to J) with no differences noted within the hypoxia or normoxia groups. **Central Illustration:** GPR39 Signaling. Upstream of GPR39, PLA_2_, is activated in hypoxia, causes greater AA release from plasma membrane. AA is then converted by 15-LO to 15-HETE, which then acts as a GPR39 agonist and activates several G-proteins (Gα_q_, Gα_s_, and Gα_12/13_), initiating downstream signaling. Gα_q_ activates PKC, increasing levels of iNOS. Gα_12/13_ stimulation results in activation of the PI3K-AKT pathway that influences expressions of VEGFA, eNOS, HIF-1_α_, and HIF-2_α_. HIF-1_α_, in turn, binds to the endothelin-1 promotor causing formation of endothelin-1. The PI3K-AKT pathway also activates the SHH pathway as well as COX2 resulting in prostaglandin formation. Both Gαq and Gα12/13 pathways stimulate the rho pathway with activation of ROCK. Gα_s_ stimulation results in increased CREB. All three pathways ultimately lead to cell proliferation and growth through transcription factors within the nucleus. The asterisk in red indicates mRNAs that are not increased in GPR39 KO compared to WT mice after 4 weeks of hypoxia. Symbol p indicates phosphorylated protein. Upward facing blue arrows show increases measured in hypoxia versus normoxia in all animals, while upward facing red arrow shows increase in WT hypoxia animals only. 15-HETE=15-hydroxyeicosatetraenoic acid; 15-LOX= 15-lipooxygenase; AA=arachidonic acid; cAMP=cyclic adenosine monophosphate; CREB=cAMP response element binding protein; COX2=cyclooxygenase 2; DAG=1,2 diacylglycerol; eNOS=endothelial nitric oxide; ERK= extracellular signal related kinase; GLI1=GLI family zinc finger 1; HIF-1α=hypoxia-induced factor 1 HIF-2α=hypoxia-induced factor 2α; iNOS=inducible nitric oxide; IP3=inositol 1,4,5 triphosphate; MAPK=mitogen-activated protein kinase; MTOR=mechanistic target for rafamycin; NFκB=nuclear factor κB; PGE= prostaglandin E; PGI=prostaglandin I PIP2 =phosphatidylinositol 4,5 biphosphate; PIP3= phosphatidylinositol (3,4,5)-trisphosphate; PKC**=**protein kinase C; PLA_2_=phospholipase A_2;_ ROCK=Rho-associated coiled-coil kinase; SHH=sonic hedgehog; SMO= smoothened; SUFU=suppressor of fused; VEGFA=vascular endothelial growth factor.

## Discussion

The new information from this study is that GPR39 is present in primary lung VSMCs and pericytes isolated from WT mice; GPR39 is also expressed in pulmonary arterioles of WT but not GPR39 KO mice; and GPR39 is also abundant in the whole lung tissue of WT mice as determined by qPCR. Genetic deletion of GPR39, the target receptor for 15-HETE, prevents hypoxia induced PAH that is associated with lesser pulmonary arterial remodeling and higher capillary density compared to littermate WT mice exposed to the same period (4 weeks) of hypoxia. mRNA and protein analysis of lung tissue indicates that GPR39 deletion abolishes a significant portion of abnormal signaling seen during hypoxia. These results suggest that GPR39 is upstream of many aberrant pathways that are activated in PAH. Hence, pharmacologically targeting GPR39 may offer a novel avenue for treatment of PAH.

### 15-HETE and GPR39

We previously reported that 15-HETE is the endogenous agonist for GPR39, which is present, among other cells, in arteriolar VSMCs^17^ and pericytes of the mouse heart.^18^ 15-HETE activates GPR39 at nanomolar concentration in the presence of physiological concentrations of Zn^++^ ^17,18^ that acts as an allosteric modulator.^19,20^ In our previous experiments, GPR39 activation by 15-HETE led to increased cytosolic Ca^++^ levels in VSMCs and pericytes^17,18^ suggesting Gα_q_ activation. Gα_q_stimulation also stimulates protein kinase C (PKC), which promotes iNOS production^36^ and alters gene expression through the mitogen-activated protein kinase (MAPK)-ERK pathway, increasing cell growth and proliferation.^37^ In our previous experiments, 15-HETE also caused ERK phosphorylation^17^ indicating stimulation of either Gα_q_ or Gα_12/13_ or both.^38^

GPR39 exhibits significant constitutive activity via its Gα_q_ and Gα_12/13_ subunits.^19,20^ GPR39 activation by synthetic analogs has been reported to also stimulate Gα_s_ causing CREB transcription induction^10,21^ leading to altered expression of genes associated with apoptosis and cell death.^39^ GPR39 activation has also be reported to stimulate Gα_12/13,_ which induces serum response element-mediated transcription that affects cell survival.^38^ Activated Gα_12/13_ stimulates the AKT-PI3K pathway, thus affecting several nuclear transcriptional factors leading to cell growth and proliferation as well as reduced apoptosis.^23,24^ The AKT-PI3K pathway acts as a regulator of HIF-1α and HIF-2 ^25^, eNOS^26^, and COX2.^27^ The AKT-PI3K axis also influences VEGFA regulation.^28^ HIF-1α, in turn, binds to endothelin-1 promotor causing formation of endothelin-1.^29^ Additionally, the AKT-PI3K activates the SHH pathway.^30,31^ Furthermore, Gα_12/13_ activation through the Rho kinase pathway also contributes to cell proliferation and growth.^38^ Activation of all these pathways have been reported in PAH.^39–41^ Because GPR39 is upstream of all these pathways, its deletion could inhibit them (central illustration), implying that pharmacological blockade of GPR39 may offer a novel approach for attenuating PAH pathophysiology.

AA and 15-HETE are upstream of GPR39 and hence their levels would not be expected to change with GPR39 deletion. To our knowledge ours is the first study to show that lung tissue AA levels are elevated in hypoxia and that 15-HETE/AA ratio is not different between hypoxia and normoxia, indicating that increased AA tissue levels may be one reason for elevated levels of 15-HETE in PAH rather than 15-LO activation alone^4,5^. Increased AA tissue levels may result from death of inflammatory cells that migrated into the pulmonary arterioles and other vascular cells early during hypoxia.^43^ Hypoxia also upregulates Phospholipase A_2_ (PLA_2_), causing greater release of AA into tissue (central illustration).^44^

As evidence of activation of Gα_q_, Gα_s_, and Gα_12/13_ pathways after 4 weeks of hypoxia, mRNA expression of ATK, CREB, ERK, and ROCK2 were elevated in WT but not GPR39 KO mice. Phosphorylated AKT levels were also elevated from hypoxia in WT but not GPR39 KO mice. That genetic deletion of GPR39 abolished stimulation of these pathways during hypoxia indicates that these pathways are downstream of GPR39. In further support of this contention, hypoxia induced mRNA and protein expression even further downstream of AKT-PI3K were blocked by GPR39 deletion: in comparison to WT hypoxia mice, GPR39 KO hypoxia mice showed no increase in mRNA expression of iNOS, HIF-1α, HIF-2α, VEGFA, and COX2 with its prostaglandin products, as well as factors involved in the SHH pathway (central illustration). In terms of protein levels, hypoxic WT animals exhibited increase in SHH, and both hypoxic WT and GPR39 KO animals exhibited increase in eNOS compared to normoxia controls. Endothelin-1, SMO, SUFU, and GLI1 expressions were downregulated in both hypoxia groups. Otherwise, there was no difference in protein expression of prostaglandin-E, COX-2, or iNOS between hypoxia and normoxia in both WT and GPR39 KO mice.

Discordance between mRNA and protein levels is not unusual. Concordance is seen, at best, in <50% of cases.^45,46^ Hypoxia itself can also cause global translation suppression despite mRNA presence.^47–49^ Further, CREB and ERK 1/2 and some hypoxia induced proteins such as H1F-1_α_, HIF-2_α_, VEGFA, and endothelin-1 may have been expressed in the early stages of hypoxia and subsided by 4 weeks.^50–52^ Despite GPR39 deletion, eNOS levels were elevated during hypoxia similar to that seen in WT mice. Whereas, the AKT-PI3k pathway is a major regulator of eNOS, other factors such as increased shear stress during hypoxia could increase eNOS levels.^53^ Interestingly, in GPR39 KO mice but not WT mice, increased expression of eNOS was associated with increased capillary density. Whether the increased pericyte density in GPR39 KO mice played a role in this finding is unclear.^54^

It was recently reported that GPR39 inhibits the SHH pathway and its deletion allows activation of the canonical pathway leading to upregulation of SHH, SMO, SUFU, and GLI1.^55^ In a chronic limb ischemia model in diabetic mice, blood flow was reported to increase in GPR39 KO mice compared to controls. We found no evidence of SHH stimulation in GPR39 KO mice, either during normoxia or hypoxia. Rather, it was only WT hypoxia mice that exhibited elevated expression of SHH with downregulation of SMO, SUFU, or GLI1 although their mRNA levels were elevated. As previously reported^56–58^, it is possible that SHH acted on the pulmonary arterial VSMCs through the noncanonical pathway, resulting in pulmonary vascular remodeling seen in our study.

### Vascular effects of GPR39 Inhibition in PAH

Our results clearly show that, unlike WT mice, GPR39 KO mice subjected to 4 weeks of hypoxia do not develop PAH. There is no evidence of increased RV systolic wall stress or RV diastolic dysfunction (normal RVEDP). Yet, there is modest pulmonary arteriolar remodeling in these animals that is coupled with increased capillary density.

Pulmonary capillary hemodynamics can be influenced by pulmonary arterial pressure. In PAH, the elevated pulmonary artery pressures can be transmitted distally to capillaries. To prevent this from happening, pre-capillary sphincters can contract reducing both capillary dimension and density (capillary blood volume).^59–61^ This protective mechanism may contribute to the hypoxemia noted in PAH^62^, at the same time preventing the occurrence of pulmonary edema. Whereas we did not see an appreciable decrease in capillary density in hypoxic compared to normoxic WT mice, we did not measure capillary diameter in this study. A small change in capillary diameter will affect microvascular resistance more than a similar change in capillary number.^63,64^ In a previous study of myocardial ischemia we noted a smaller capillary diameter in WT compared to GPR39 KO mice.^18^

Capillaries in the lungs are placed in parallel and so an increase in capillary number (and diameter) lessens total vascular resistance.^63,64^ Thus, increased capillary density may explain GPR39 KO mice’s lack of developing PAH despite modest remodeling of pulmonary arterioles. In the normal lung, because of high pulmonary vascular compliance, addition or deletion of even a large number of capillaries does not affect pulmonary vascular resistance. However, when pulmonary circulatory compliance is restricted from pulmonary arterial thickening, capillary density becomes a far more important contributor to total pulmonary vascular resistance. Increased capillary density could, therefore, represent a compensatory mechanism.

15-HETE itself has been reported to induce angiogenesis not only in PAH^6^ but also other conditions such as cancer^65^ and diabetic retinopathy.^66^ Unlike intussusceptive angiogenesis noted in our study, that reduces microvascular resistance by increasing capillary density, 15-HETE induces sprouting, proliferative, maladaptive angiogenesis causing new vessel formation without beneficial hemodynamic effects, and instead may lead to leaky blood vessels, edema, and hemorrhage. GPR39 blockade may thus prevent this type of unwanted angiogenesis.

15-HETE causes VSMC contraction by increasing intracellular Ca^++^.^12,17^ VSMCs or pericytes isolated from GPR39 KO mice do not demonstrate this effect.^17,18^ This mechanism of vasoconstriction, which might contribute to high pulmonary artery pressure is abolished with GPR39 deletion. 15-HETE also inhibits K^+^ channels, contributing to vasoconstriction.^13–15^ This effect is likely downstream of GPR39, and its blockade could be an additional mechanism that ameliorates the vasoconstrictor effects of 15-HETE in PAH.

Pericyte coverage around distal pulmonary arterioles is known to increase in PAH^9,67,68^, which may indicate an increased role of pericytes in PAH. Pericytes also surround capillaries and control local capillary flow.^59,60^ Here we show that at the capillary level GPR39 KO mice subjected to either normoxia or hypoxia have markedly increased pericyte density compared to WT mice, most likely a developmental feature caused by GPR39 ablation. Inhibition of PI3K and Rho, both downstream of GPR39, have been shown to affect pericyte maturation and function.^69,70^ Because of pericyte-capillary interactions, abundant pericytes in the GPR39 KO mice could have contributed to increases capillary density in these mice during hypoxia.

### Limitations of our Study

We selected the mouse rather than the more commonly used rat model of PAH because of the availability of GPR39 KO animal and its phenotypical as well as functional characterization in our laboratory. We did not see several reported abnormalities in hypoxia models, such as upregulation of VEGFA, COX-2, endothelin-1, and iNOS, most likely because the increases had subsided at 4 weeks or because of hypoxia-induced global suppression of translation. The decrease in CREB, HIF-1_α_, HIF-2_α_ and endothelin-1 after 4 weeks of hypoxia despite increased mRNA expression could be explained by increased protein degradation and/or other compensatory mechanisms.

Whereas we noted abolishment of many abnormal signaling pathways reported in PAH, we nonetheless saw some pulmonary arterial thickening and remodeling. It is possible that factors not measured in our study may be responsible for this finding. Transforming growth factor β (TGFβ) and Bone morphogenetic protein receptor type II (BMPR2) and their interactions, are possible additional targets, among others, that may affect tissue growth and proliferation.^71,72^

### Therapeutic Implications of GPR39 Blockade in PAH

PAH is a devastating disease with very poor outcome.^1^ Current standard treatments address metabolic and structural abnormalities that are end results of abnormal signaling in the lung.^1–3^ Although 15-HETE has been reported to disrupt many of these pathways^4^, the receptor through which 15-HETE acts was unknown. We reported that 15-HETE is the endogenous agonist for GPR39. Here we show that genetic deletion of GPR39 abolishes many abnormal signaling pathways associated with hypoxia, significantly attenuating pulmonary arterial remodeling and increasing capillary density, thus preventing development of PAH. Our results suggest that GPR39 plays an important role in the pathogenesis of PAH and should be targeted pharmacologically for the treatment of PAH.

## Abbreviations and Acronyms

15-HETE: 15-hydroxyeicosatetraenoic acid
15-LOX: 15-lipooxygenase
cAMP: cyclic adenosine monophosphate
CREB: cAMP response element binding protein
COX2: cyclooxygenase 2
eNOS: endothelial nitric oxide
ERK: extracellular signal related kinase
GLI1: GLI family zinc finger 1
HIF-1α: hypoxia-induced factor 1
HIF-2α: hypoxia-induced factor 2α
ICC: immunocytochemistry
IHC: immunohistochemistry
iNOS: inducible nitric oxide
IP3: inositol 1,4,5 triphosphate
MAPK: mitogen-activated protein kinase
PGE: prostaglandin E
PGI: prostaglandin I
PLA_2_: phospholipase A_2_
qPCR: quantitative polymerase chain reaction
ROCK: Rho-associated coiled-coil kinase
SHH: sonic hedgehog
SMO: smoothened
SUFU: suppressor of fused
VEGFA: vascular endothelial growth factor A

## Author Contributions

SK and CM developed the concept and design for the studies and performed the experiments. SK performed hemodynamic and image analysis. CM performed and analyzed all western blot data. MP isolated VSMCs and pericytes, LL performed IHC, PC performed ICC and AT performed qPCR. Figures were created by CM, AT, PC, AT and SK. All authors contributed to drafting their portions of the manuscript with SK drafting the final version.

## Acknowledgements

Metabolomic analysis was performed by the Bioanalytical Shared Resource/ Pharmacokinetics Core Facility (Research Resource Identifier (RRID): SCR_009963), which is supported in part by the University Shared Resource Program at Oregon Health and Sciences University. We are thankful to the integrated genomic laboratory for use of qPCR equipment. It receives its support from the National Cancer Institute Center Support Grant P30CA069533 to the Knight Cancer Institute. The research reported in this publication used computational infrastructure supported by the Office of Research Infrastructure Programs, Office of the Director, of the National Institutes of Health under Award Number S10OD034224. The content is solely the responsibility of the authors and does not necessarily represent the official views of the National Institutes of Health.

We thank Elizabeth Le, MD, for creating the central illustration in Biorender.

## Central Illustration

GPR39 Signaling.

## Central Illustration

GPR39 Signaling.

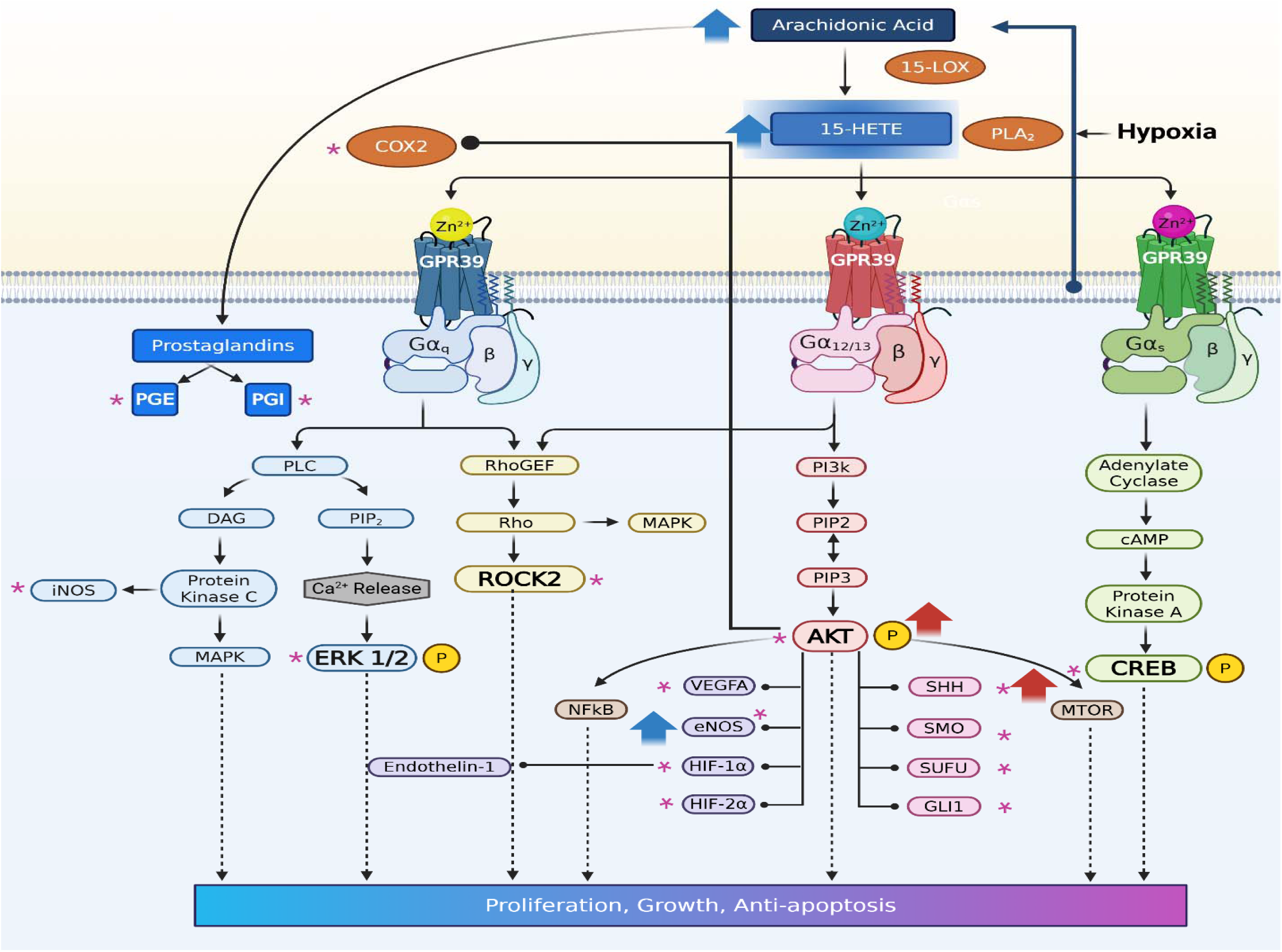

